# *Quantum CART* (*qCART*), a *piggyBac-based* system for development and production of virus-free multiplex CAR-T cell therapy

**DOI:** 10.1101/2022.05.03.490469

**Authors:** Yi-Chun Chen, Wei-Kai Hua, Jeff C. Hsu, Peter S. Chang, Kuo-Lan Karen Wen, Yi-Wun Huang, Jui-Cheng Tsai, Yi-Hsin Kao, Pei-Hua Wu, Po-Nan Wang, Ke-Fan Chen, Wan-Ting Liao, Sareina Chiung-Yuan Wu

## Abstract

Chimeric antigen receptor T (CAR-T) cell therapy has the potential to transform cancer treatment. However, CAR-T therapy application is currently limited to certain types of relapsed/refractory B cell lymphomas. To unlock the full potential of CAR-T therapy, technologic breakthroughs will be needed in multiple areas, including optimization of autologous CAR-T development, shortening the innovation cycle, and further manufacturing advancement of next-generation CAR-T therapies. Here, we established a simple and robust virus-free multiplex *Quantum CART^™^* system that seamlessly and synergistically integrates four platforms: 1. *GTailor^™^* for rapid identification of lead CAR construct design, 2. *Quantum Nufect^™^* for effective but gentle electroporation-based gene delivery, 3. *Quantum pBac^™^*, featuring a virus-free transposon-based vector with large payload capacity and integration profile similar to retrovirus, and 4. *iCellar^™^* for robust and high-quality CAR^+^ T memory stem cell expansion. This robust, virus-free multiplex *Quantum CART^™^* system is expected to unleash the full potential of CAR-T therapy for treating diseases.

## Introduction

In recent decades, immunotherapy involving T lymphocytes capable of antigen-specific recognition leading to cytotoxic activity and persistence, have taken center stage in tumor eradication.^1,2^ Among T cell-based therapies, chimeric antigen receptor T (CAR-T) therapies that combine the specificity of antibody recognition and T cell activating functions into a single protein receptor, have shown considerable potential to advance cancer treatment.^3 4^

A successful CAR-T therapy requires not only tumor recognition, but also sufficient cell expansion, persistence, and delivery of effector functions. Second generation CAR-T cells that incorporate CD28 or 4-1BB in the CAR design have demonstrated therapeutic efficacy against hematological malignancies.^5^ However, successful treatment against solid tumors remains a challenge due to the hostile immunosuppressive tumor microenvironment (TME) that inhibits the infiltration, survival, and cytotoxic activity of CAR-T lymphocytes.^6–8^ One strategy to overcome this challenge is to engineer CAR-T cells to express additional genes that enhance CAR-T functions, so called armored CAR-T cells.^9^ Efficacy of armored CAR-T therapies against solid tumors in preclinical studies has been well-documented, and clinical trials are currently ongoing.^10–15^ However, the repertoire of genes that can be included in armored CAR-T therapies has been severely restricted by the limited cargo capacity of viral vectors. Another strategy is to enrich for specific T cell subsets that possess superior capability to persist and attack cancer. Ever since Gattinoni *et al*. described memory stem T cells (T_SCM_),^16^ several studies have demonstrated T_SCM_ to be critical in facilitating successful adoptive T cell therapy.^17–20^

Currently, retroviral and lentiviral vectors are the most common vehicles used for CAR-T engineering.^21,22^ However, virus-based CAR-T therapies have potential risk of genotoxicity and are unsuitable for multiplex CAR-T production due to limited gene cargo capacity. Furthermore, the high cost of virus-based CAR-T therapies prevent widespread patient accessibility. Non-virus-based CAR-T therapy is a promising alternative. However, until more recently its development has been hampered by low gene transfer efficiency and difficulty to efficiently expand quality engineered T cells for clinical application.^23^

In this study, we describe virus-free *Quantum Engine^™^*, a cell engineering and production system composed of four platforms: *GTailor^™^, Quantum Nufect^™^, Quantum pBac^™^*, and *iCellar^™^*. We demonstrate that *qCART^™^* (a *Quantum Engine*^™^ for CAR-T production) can generate multiplex CAR-T cells that express genes of interest (GOIs) with various targeting specificities and sizes. The engineered CAR-T cells consistently exhibit desired features, including high T_SCM_, high expansion capacity, and robust anti-tumor efficacy.

Given that *qCART^™^* can shorten both the development and manufacturing timeline of virus-free multiplex CAR-T therapy, *qCART^™^* represents a technological breakthrough that will be expected to unlock the full potential of CAR-T therapies and facilitate widespread patient access.

## Results

### *GTailor^™^* facilitates screening of therapeutic gene constructs to identify lead candidates for preclinical development

Since identification of CAR-T lead candidates is laborious, time-consuming and requires multiple rounds of screening involving designing and construction of gene constructs and subsequent *in vitro* and/or *in vivo* testing and validation, we developed the *GTailor^™^* platform to streamline these processes for efficient lead candidate identification. This platform consists of the following technologies: (1) a library of functional modules for rapid establishment of a therapeutic construct, (2) a simple, time-effective and economical cell engineering process for building a multiplex CAR-T cell library, (3) a high throughput *in vitro* cytotoxicity assay system for identifying potent, persistent CAR-T cells that possess high On-Target, and minimal Off-Tumor toxicities. A sample screening process for B-cell maturation antigen (BCMA) CAR-T cells is depicted in Supplementary Fig. S1, and the results are presented in Table 1. As shown, all five groups of CAR-T cells performed similarly and satisfactorily in terms of cell expansion (day 1 to 10-fold change), percentage of CAR^+^ cells, CD8/CD4 ratio, and percentage of T_SCM_ in both CD4 and CD8 populations (Table 1).

**Table 1.** Finding optimal BCMA CAR-T cells. Human peripheral blood mononuclear cells (PBMC) electroporated with *Quantum pBac^™^*expressing anti-BCMA CARs were analyzed for their performance, including cell expansion (Day 1 to 10 fold change), percentage of CAR^+^ cells, CD8/CD4 ratio, percentage of T_SCM_ cell subsets in CD4^+^ or CD8^+^ cells, and cytotoxicity against BCMA^+^ cells. Data shown are from four healthy donors. Results are shown as mean fold change, CD8/CD4 ratio, cytotoxic activity, or percentage of CAR^+^, CD4^+^/T_SCM_, CD8^+^/T_SCM_, or CAR^+^/PD1^+^ cells.

| Group | N<br>(independent experiments) | Day 1 to 10<br>fold change | % CAR<sup>+</sup> | CD8/CD4<br>ratio | % T<sub>SCM</sub><br>in CD4<sup>+</sup> | % T<sub>SCM</sub><br>in CD8<sup>+</sup> | % PD1<sup>+</sup><br>in CAR<sup>+</sup> | Cytotoxicity<br>against<br>BCMA<sup>+</sup> cells<br>(5:1, 48hr) |
| --- | --- | --- | --- | --- | --- | --- | --- | --- |
| aBCMA-1 | 2 | 168<br>(101, 236) | 45.8<br>(45.0, 46.6) | 1.3<br>(0.9, 1.7) | 60.3<br>(59.5, 61.1) | 74.6<br>(73.3, 75.9) | NA | 15.8<br>(14.7, 16.8) |
| aBCMA-2 | 7 | 226<br>(141~384) | 63.7<br>(55.3~74.5) | 1.5<br>(0.4~3.8) | 71.0<br>(63.3~80.6) | 76.3<br>(63.3~83.1) | 0.8<br>(0.6~1.7) | 55.3<br>(46.0~71.5) |
| aBCMA-3 | 3 | 289<br>(162~472) | 51.6<br>(33.9~73.4) | 2.7<br>(1.4~4.2) | 76.6<br>(67.6~84.7) | 82.4<br>(75.4~87.4) | 74.6<br>(49.9~90.4) | 9.8<br>(3.3~14.2) |
| aBCMA-4 | 2 | 223<br>(149, 297) | 57.8<br>(50.5, 65.1) | 3.4<br>(1.8, 5) | 68.1<br>(64.9, 71.3) | 69.5<br>(57.4, 81.6) | 6.4<br>(1.2, 11.5) | 12.5<br>(11.7, 13.2) |
| aBCMA-5 | 3 | 278<br>(187~454) | 68.3<br>(68.1~68.5) | 2.5<br>(1.9~3.7) | 73.6<br>(67.1~80.4) | 79.7<br>(71.7~85.7) | 0.6<br>(0.4~1.1) | 50.5<br>(43.2~57.5) |

However, three CAR-T groups (aBCMA-1, aBCMA-3 and aBCMA-4) exhibited markedly lower cytotoxicity against BCMA^+^ target cells. Note that the percentage of PD1^+^ CAR-T cells in the aBCMA-3 group was also markedly higher than those found in other groups. CAR-T cells with these undesired traits were tested at least twice (using T cells derived from different donors), whereas other groups were tested at least three times to confirm reproducibility. As a result, aBCMA-2 and aBCMA-5 CAR-T cells were identified as tentative candidates. These two candidate CAR-T groups were next co-cultured with a panel of BCMA-negative and BCMA-positive tumor and normal cells. As shown (Supplementary Table S1), aBCMA-5 lysed more cells expressing minimal BCMA, therefore aBCMA-2 was chosen as the candidate for pipeline development.

### *Quantum pBac^™^ (qPB)* facilitates consistent intra-cell type genome integration

We have previously demonstrated that the *qPB* system is superior compared to other *piggyBac* transposon systems.^24^ Since the integration characteristics of *qPB* is unknown, we analyzed for the first time, the integration profile of *qPB* in CAR-T cells. Additionally, we compared the integration profile with that of human embryonic kidney (HEK293) cells^25^ to determine whether there may be any inter-cell type differences. As shown in Figure 1A, integration sites (IS) were identified within all T cell chromosomes with the exception of chromosome Y. This may be due to the relatively smaller size of Y chromosome, and would be consistent with the lower number of IS found within relatively smaller chromosomes such as chromosomes 13-22 (Figure 1A and 1B). In contrast to the relatively consistent IS biodistribution pattern found in T cells derived from two separate donors, several distinct differences were found in integration profile between HEK293 and T cells. For example, in contrast to T cells, no IS were found within chromosome 16 of HEK293 (Figure 1A and 1B). Moreover, higher percentages of IS were found within chromosomes 5 and 6 of HEK293, whereas more IS were found within chromosome 19 for T cells (from both donors; Figure 1B).

**Figure 1.**
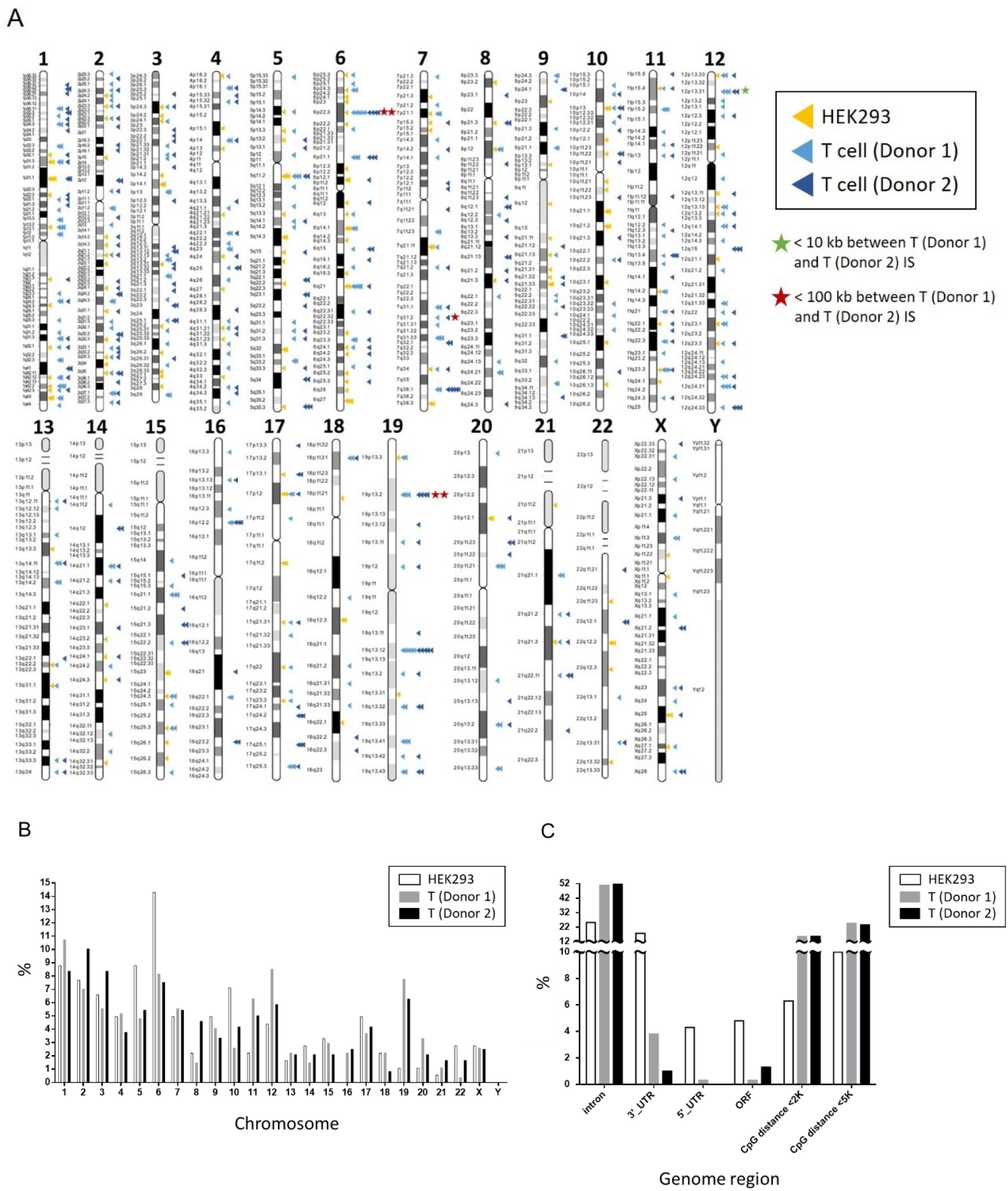
Profiling of *Quantum pBac^™^ (qPB)* genome integration. (A) A schematic depiction showing the biodistribution of *qPB* genome integration sites (IS) mapping to specific segments of the indicated HEK293 and primary T cell chromosomes. (B), (C) IS as shown in (A), data analyzed and grouped by: the indicated chromosome (B), and the indicated genome region (C). Results shown are from two females (HEK293, T donor 2) and one male (T donor 1), and results of (B) and (C) are presented as percentages of total IS.

Since *qPB* genome insertion may raise safety concerns, we next analyzed the biodistribution of IS within or near different areas of genes. A total of 344 and 300 unique IS were identified within CAR-T cell donor 1 and donor 2, respectively (Supplementary Table S2). Of note, we observed that increasing the amount of donor DNA (carrying CAR) introduced into CAR-T cells increases integrant copy number in cells.^26^ To reduce safety risks, we optimized the concentration of donor DNA such that integrant copy number per cell would be below five, as shown by the low CAR copy numbers found in T cells of both donors (3.08 and 1.68 in donors 1 and 2, respectively; Supplementary Table S2). Interestingly, remarkable conformity was found in the integration profiles of both donor T cells. For example, approximately 50% of IS (50.9% and 52% in donors 1 and donor 2, respectively) were found in the introns, while a very low percentage (0.3% and 1.3% in donors 1 and donor 2, respectively) were found in the open reading frame (ORF) of protein coding gene (Figure 1C). Historical data demonstrated that lentiviral vector with an even higher percentage (79.7%) of intronic IS^27^ is considered to be relatively safe. This suggests that *qPB* may be as safe as lentiviral vectors, although other factors including bias concerning targeting genes and transcriptional start sites/termination sties remain to be assessed. Furthermore, the lower percentage of IS found in the introns (25.2%) and higher percentage of IS found in the ORF (4.8%) of HEK293 cells support cell type-specific biodistribution of *qPB* IS. Next, we focused on IS in the CpG dinucleotide-rich regions (CpG islands) since the transcription of >50% of human genes are initiated from these regions.^28^ Similar percentages of IS were found <5 Kb from CpG islands (24.7% and 23.7% in donors 1 and donor 2, respectively) as compared to published data in cells transduced using retrovirus (32.8%) or hyperactive *piggyBac* (*hyPB*, 26.6%).^27^.

While the majority of IS were identified in genes, none of the top 10 IS of either T cell donors targeted within known cancer-associated genes (Supplementary Table S3). Importantly, some IS were found within or in proximity to cancer-associated genes, but they were either in the intronic region, in the UTR region, or upstream of these genes (Supplementary Table S4). Moreover, no IS was found in/near any of the genes (*CCND2, HMGA2, LMO2, MECOM, MN1, PRDM16*) previously reported to be associated with severe adverse events in patients.^29–32^ This biodistribution pattern of IS thereby suggests that *qPB* may be as safe as currently available randomly-integrating vector systems.

### *Quantum Nufect^™^ (qNF*) facilitates effective transfection of human primary T cells while preserving high cell viability

Electroporation is the most efficient method for delivery of virus-free vectors into therapeutic cells. However, a major bottleneck of this approach includes low viability and expansion capacity of electroporated cells. To address this matter, we developed a cell type-independent generic buffer system for electroporation called *Quantum Nufect^™^ (qNF)*. We utilized Lonza’s Nucleofector^™^ to assess the effect of *qNF* on T cell transfection efficiency. We first nucleofected T cells with a pmaxGFP plasmid using *qNF*, and compared the results to those obtained using the Human T Cell Nucleofector^™^ Solution (HTCN, Lonza; Figure 2A).

**Figure 2.**
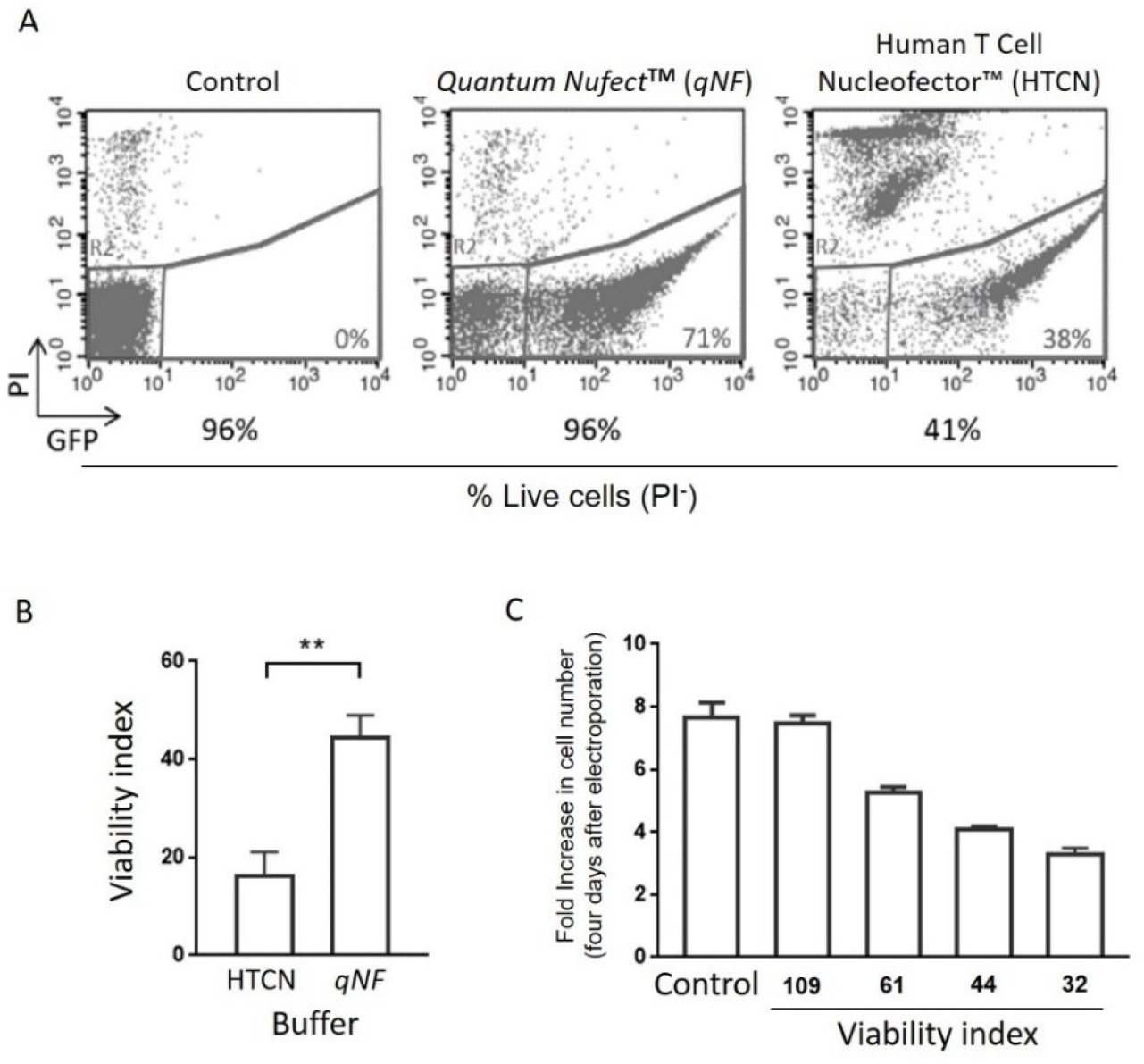
Viability of human primary T cells electroporated using *Quantum Nufect^™^ (qNF*) and compared with those electroporated using the Human T Cell Nucleofector^™^ solution. Human peripheral blood CD8^+^ T cells electroporated with pmaxGFP plasmid in either *qNF* buffer or Human T Cell Nucleofector^™^ (HTCN) solution. One day after electroporation, transfected cells were analyzed for (A) GFP and PI staining, and (B) viability index. (C) Human peripheral blood T cells electroporated under conditions that result in different levels of post-electroporation viability were analyzed for their viability indices plotted versus fold increase in cell number after four days of culture. Results are shown as percentage of cells positive or not positive for GFP and/or PI staining (A), mean viability index (B, C) and the mean fold increase in number of cells (C). ** *p* < 0.01. N = 3 (B, C, triplicates).

We show in a representative experiment, that unlike the low viability (41%) seen in HTCN-transfected T cells, *qNF* preserved the viability of a high percentage (96%) of cells at a level comparable to non-nucleofected control cells. These results are consistent with a significantly higher viability index in *qNF*-nucleofected cells (Figure 2B). We next examined the relationship between cell viability and expansion capacity of T cells transfected with pmaxGFP using *qNF*. As shown in Figure 2C, low initial viability resulted in low subsequent expansion after three additional days of culture, suggesting that cell viability following nucleofection is positively associated with cell expansion capacity.

Next, we assessed the effect of *qNF* on CAR transfection efficiency by nucleofecting cells with *qPB* carrying a transcript expressing a tandem CD20/CD19 CAR, and iCasp9, using *qNF* or HTCN. As shown in Figure 3A, significantly more total live cells were recovered from the *qNF* group (8.9E5 ± 9.3E4) compared with those recovered from the HTCN group (3.2E5 ± 1.0E5) one day after nucleofection. While a significantly higher percentage of live cells of the HTCN group were CAR^+^, nearly twice as many CAR^+^ T cells were recovered using *qNF* (2.0E5 ± 3.7E4) compared to that using HTCN (1.2E5 ± 3.8E4; Figure 3A).

**Figure 3.**
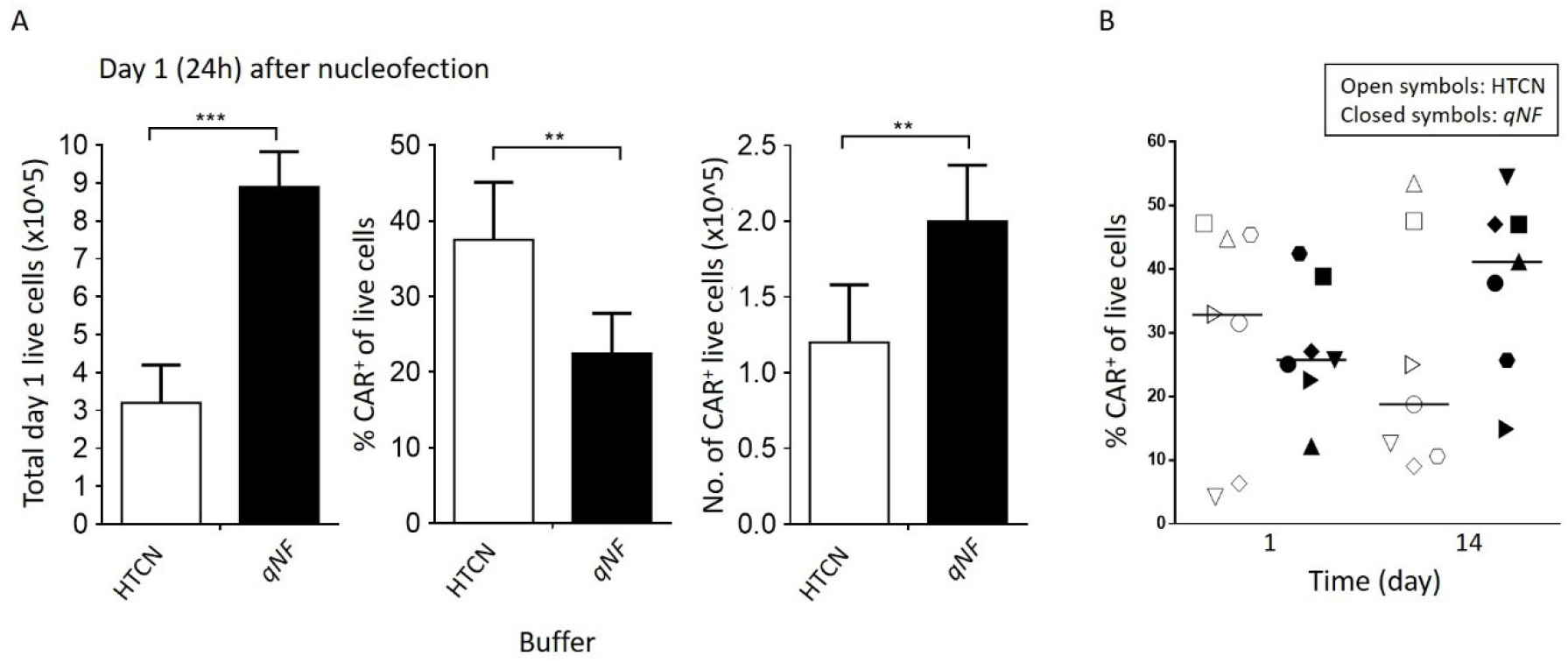
Human primary T cells electroporated using *Quantum Nufect^™^ (qNF*) buffer system produces high yield of CAR^+^ T cells. Human peripheral blood T cells electroporated with *Quantum pBac^™^* expressing CAR in either *qNF* buffer or Human T Cell Nucleofector^™^ (HTCN) solution were harvested one day after electroporation and analyzed for (A) CAR and PI staining, and yield calculation. (B) The percentages of CAR^+^ cells were analyzed on day 1 and day 14. Results are shown in (B) as median percentage of CAR^+^ live cells. Yield is calculated as No. of CAR^+^ live cells, using the formula: (Total day 1 live cells) × (percentage of CAR^+^ live cells). ** *p* < 0.01, *** *p* < 0.001. N = 3 (A, triplicates), N = 7 donors (B).

Next, we addressed whether the greater transfection rate seen in HTCN-transfected cells would result in higher percentage of CAR^+^ cells in the final CAR-T product. We compared CAR transfection efficiency using *qNF* vs HTCN in cells obtained from seven healthy donors. As shown in Figure 3B, despite an initial lower percentage of CAR^+^ T cells one day after nucleofection using *qNF*, there was an overall increase in CAR^+^ cells after 14 days of culture. This is in contrast to an overall decrease in percentage of CAR^+^ cells seen in T cells nucleofected using HTCN. In fact, in cells from five out of seven donors, the slope of increase in percentage of CAR^+^ T cells following nucleofection was steeper in *qNF* than in HTCN-nucleofected cells (Supplementary Fig. S2). Moreover, in the remaining two donors, the slope of decrease in percentage of CAR^+^ T cells following nucleofection was the same or less steep in *qNF* than in HTCN-nucleofected cells.

These evidence suggest that an initial high percentage of CAR^+^ cells will not always lead to high CAR^+^ cell enrichment at harvest, and that *qNF* is superior and more reliable in producing and enriching for CAR^+^ T cells than HTCN.

### *iCellar^™^* robustly enriches and expands CAR-T cells while maintaining cell stemness (T_SCM_)

We have previously shown that *qPB* can be advanced towards clinical application in CAR-T therapy.^24^ However, even in the presence of artificial antigen presenting cells (aAPC), T cells from some donors still failed to sufficiently expand. To resolve this issue, we developed a CAR-T cell culture supplement named *Quantum Booster^™^ (qBT*), which in conjunction with aAPC are both components of *iCellar^™^*. We analyzed the effect of *iCellar^™^* on performance of CAR-T cells by determining whether different components of *iCellar^™^*, namely aAPC, *qBT*, or their combination (aAPC+*qBT*), affect the characteristics of CAR-T cells produced. As shown in Figure 4, *qBT* increased the expansion of CAR-T cells (Figure 4A), as well as significantly enriched for CAR^+^ T cells (Figure 4B).

**Figure 4.**
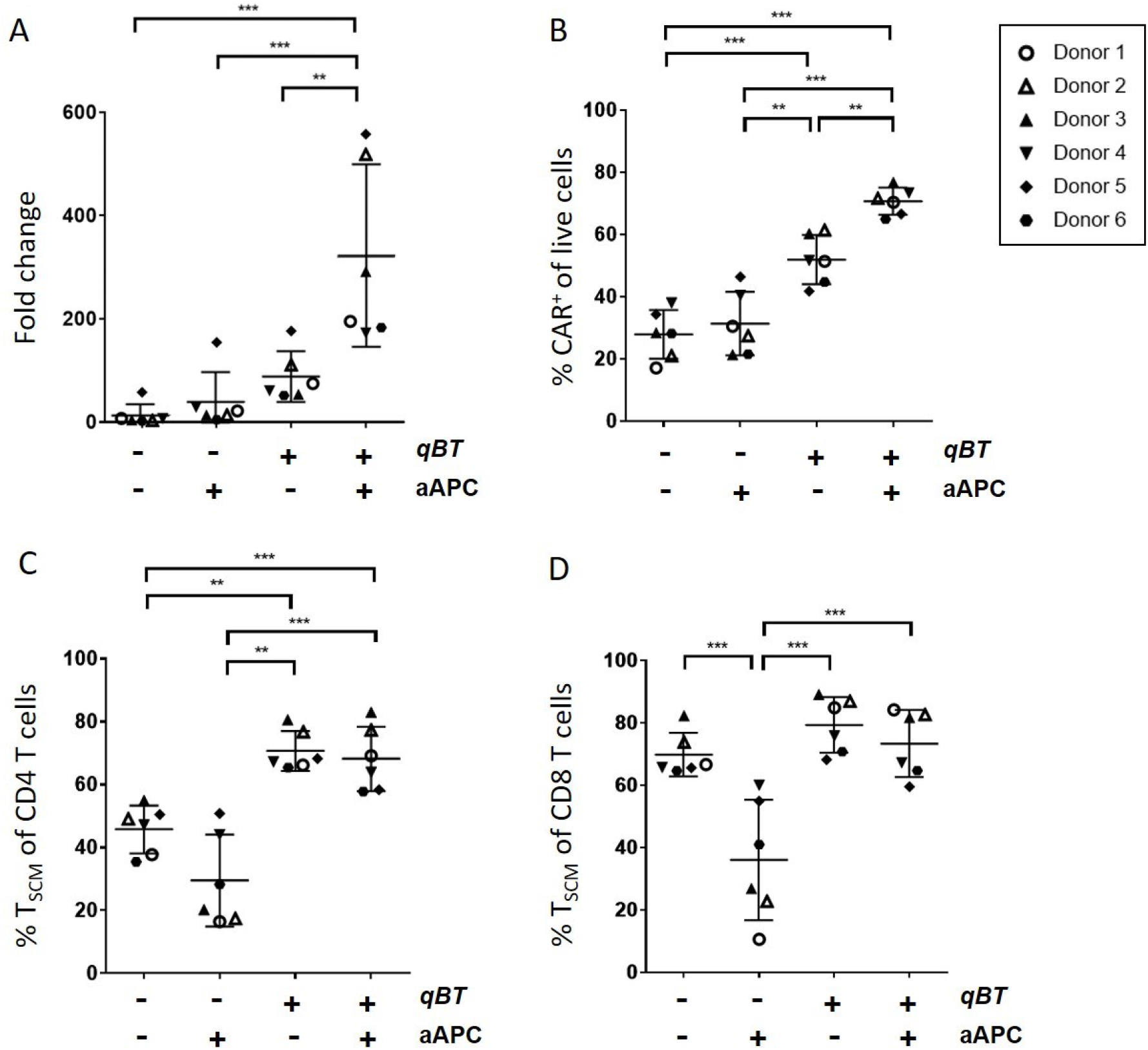
The effect of different *iCellar^™^* combinations on characteristics of CAR-T cells. Human peripheral blood mononuclear cells (PBMC) electroporated with *Quantum pBac^™^* expressing CAR and cultured for 10 days in the presence or absence of aAPC and/or *Quantum Booster^™^ (qBT*) were harvested and assessed for (A) cell expansion fold change, (B) percentage CAR^+^ of live cells (CD3^+^ PI^-^, >90%), and percentage of T_SCM_ cell subsets in (C) CD4^+^ or (D) CD8^+^ cells. Data shown are from six healthy donors. Horizontal lines represent the mean and s.e.m. fold change (A), mean and s.e.m. percentage of CAR^+^ cells (B), and mean and s.e.m. percentage of T_SCM_ cell subsets (C and D). * *p* < 0.05, ** *p* < 0.01, *** *p* < 0.001.

On the other hand, while aAPC alone did not significantly affect expansion or percentage of CAR^+^ T cells, aAPC+*qBT* significantly enhanced both of these parameters (Figure 4A and 4B). Moreover, while either *qBT* or aAPC+*qBT* increased the percentage of T_SCM_ cells over the control group (Figure 4C and 4D), aAPC alone markedly decreased the percentage of T_SCM_ cells. This effect, which reached statistical significance in the CD8^+^ population (Figure 4D) suggests that aAPC promotes cell maturation. Furthermore, aAPC+*qBT* significantly blocked the reduction in percentage of T_SCM_ cells (Figure 4C and 4D), suggesting that *qBT* maintains cells in a less differentiated state (i.e. T_SCM_). These results demonstrate that *qBT* of *iCellar^™^* markedly enhance the expansion capacity of CAR-T cells while preserving and/or even enhancing CAR-T quality.

### Effective tumor clearance by *qCART^™^*-produced CAR-T cells in Raji-bearing immunodeficient mice

To further determine the effect of aAPC on the functionality of *qCART^™^*-produced CAR-T cells, we first assessed its effect on the performance of CAR-T cells produced from four healthy donors (Supplementary Table S5). We found similar and consistent performance in cells from all four donors, and assessed cytotoxic activities of T cells from donor A. As shown in Figure 5A, CAR-T cells effectively lysed Raji-GFP/Luc cells.

**Figure 5.**
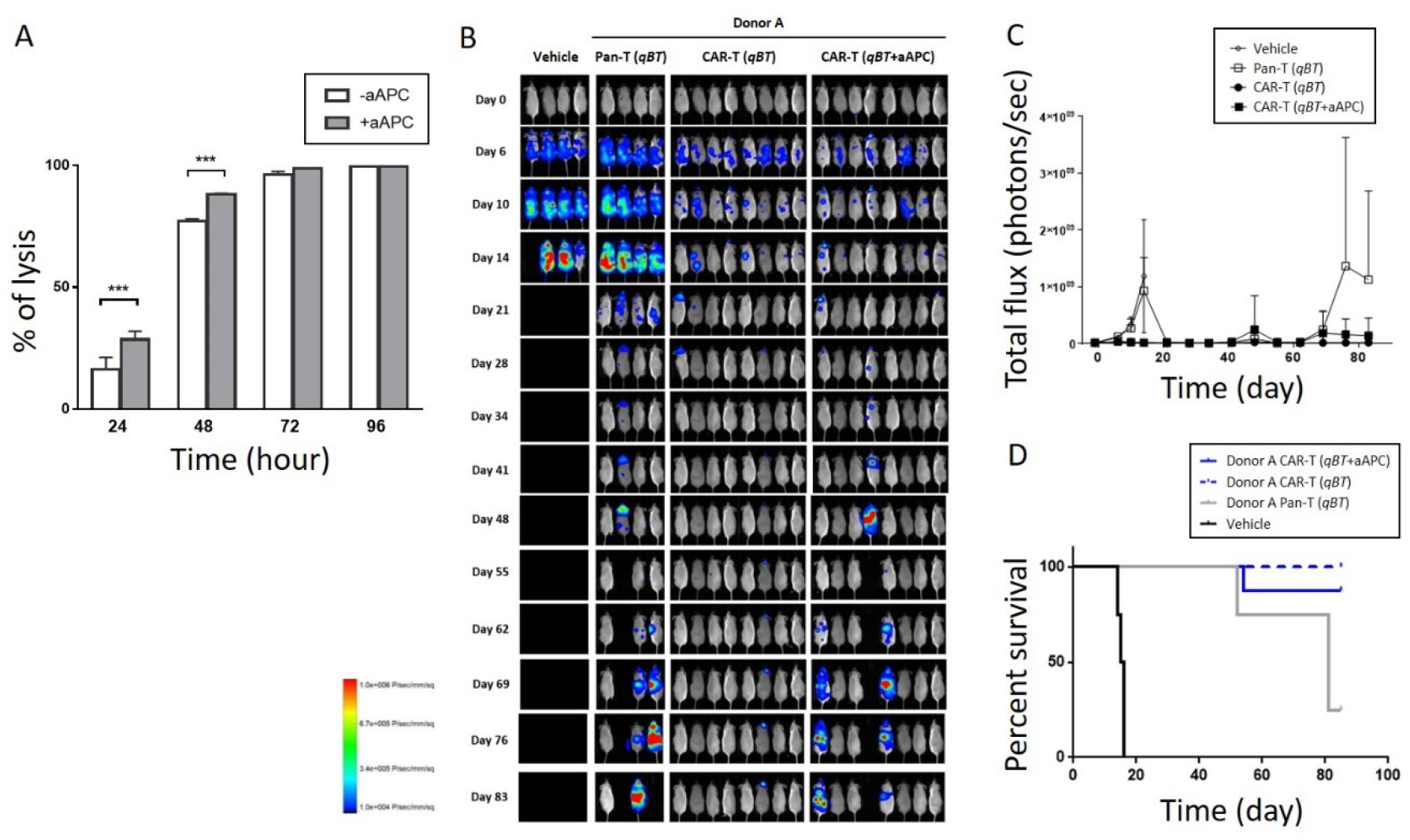
*In vitro* and *in vivo* functional characterization of CAR-T cells produced using the *qCART^™^* system. (A) *In vitro* cytotoxicity of CAR-T cells (E:T ratio of 1:1) with or without pre-incubation with aAPC, on Raji-GFP/Luc target cells. (B) *In vivo* cytotoxicity of CAR-T cells with or without pre-incubation with aAPC in Raji-GFP/Luc-bearing immunodeficient mice. Fluorescence intensity values (C) and survival curves (D) of mice from (B) plotted against time. Results shown are from four to eight mice/group. T cells were obtained from a representative donor. Vehicle and Pan-T cells (non-engineered T cells) were used as controls. *Quantum Booster^™^ (qBT*) was present in all cell culture conditions. *** *p* < 0.001.

Of note, CAR-T cells expanded with aAPC exhibited significantly increased killing efficacy at earlier (24h and 48h) time points. In contrast to vehicle and pan-T-injected mice, Raji-bearing mice injected with CAR-T cells (*qBT* or *qBT*+aAPC) killed markedly more Raji cells (Figure 5B, 5C), despite the significantly higher plasma IFN-γ levels in pan-T-injected mice (Supplementary Fig. S3A, see day 15). The superior antitumor efficacy of CAR-T cells either with *qBT* or *qBT*+aAPC was also shown by the prolonged survival of these mice compared with those injected with vehicle or pan-T cells (Figure 5D). This observation is consistent with the absence on day 85, of detectable Raji cells remaining in the blood and bone marrow of CAR-T treated mice (*qBT* or *qBT*+aAPC; Supplementary Fig. S3C), despite the continued presence of tumor in the liver and/or ovaries of tumor-bearing mice (Figure 5B, Day 83, CAR-T (*qBT*+aAPC); verified by surgery). Interestingly, we observed tumor relapse in mice that had been injected with CAR-T cells cultured with *qBT*+aAPC, but not with *qBT* alone. We also observed that in one mouse (first from the left in the “CAR-T (*qBT*)” panel) tumor relapsed by day 21. However, complete tumor remission was again observed by day 34 with no further evidence of relapse up end of experiment (Figure 5B). This observation is corroborated by the expected higher number of circulating CAR^+^ T cells (due to higher CAR copy number in blood) found in CAR-T cell-treated mice on day 71 vs day 43 (Supplementary Fig. S3B), and suggests that the increased CAR^+^ T cells may contribute to tumor clearance.

### *qCART^™^* produces CAR-T cells of different targeting specificity and gene of interest sizes with expansion capacity capable of reaching clinical scale production

To determine whether CAR-T cells with CAR genes expressing binder(s) against various antigens and/or transgene of various sizes could be effectively generated, we nucleofected *qPB* carrying various genes of interest (GOI) into T cells of healthy donors and analyzed the performance of CAR-T cells. Using the perfusion setup, we observed that increasing the transgene size decreased the percentage and expansion capacity of CAR^+^ cells (Supplementary Table S6). On the other hand, using the G-Rex setup, even though we still observed an inverse association between percentage of CAR^+^ cells and GOI sizes, the difference in CAR^+^ cell expansion appeared to be more consistent and much less affected by GOI size (Table 2).

**Table 2.** Consistent performance of CAR-T cell products produced by *qCART^™^*.

| Pipeline (Multiplex) | GOI Size (Kb) | Description | % CAR<sup>+</sup> | CAR-T Expansion (Fold Increase) | CD8/CD4 Ratio | % CD8<sup>+</sup> T<sub>SCM</sub> / % CD4<sup>+</sup> T<sub>SCM</sub> in CAR<sup>+</sup> | Exhaustion Markers PD-1/ TIM3/ LAG3 (%) | Senescence Markers KLRG1/ CD57 (%) | Representative *In Vitro* Cytotoxicity (%) |
| --- | --- | --- | --- | --- | --- | --- | --- | --- | --- |
| GF-CART01 (CD20/CD19-iCasp9) | 5.2 | iCasp9 + CD20/CD19 | 31.9~49.9 (n=6) | 158~512 | 5 | 73.6~94.7/ 65.1~82.5 | 0.4~2.0/ 0.0~5.3/ 0.9~7.0 | 2.2~14.6/ 0.0~0.7 | 32.7~92.8 (E/T=1, 48 hrs) |
| GF-CART02 (Dual CAR-iCasp9) Multiple Myeloma | 3.4 | CAR (binder 1) | 55.3~74.5 (n=7) | 141~383 | 0.4~3.8 | 63.3~83.1/ 63.3~80.6 | 0.2~1.7/ 0.5~7.6/ 0.6~3.4 | 0.4~7.9/ 0.2~10.8 | 41.6~71.3 (E/T=5, 48 hrs) |
|  | 3.4 | CAR (binder 2) | 44.4~88.7 (n=4) | 163~535 | 1~3.5 | 74.4~79.2/ 53.3~71.3 | 0.2~3.6/ 0.8~4.2/ 0.9~4.5 | 7.8~24.5/ 0.1~1.2 | 19.3~39.1 (E/T=5, 48 hrs) |
|  | 5.4 | iCasp9-CAR (binder 1+2) | 48.4, 58.5 (n=2) | 134, 243 | 1.8, 1.8 | 68.8, 72.2/ 75.6, 75.7 | 0.2, 0.4/ 0.7, 1.3/ 1.6, 3.1 | 0.5, 0.6/ 0.2, 0.4 | 34.8, 68.8 (E/T=5, 48 hrs) |
| GF-CART03 Multiple Solid Tumors | 7.3 | iCasp9-CAR Modulator 1 + Modulator 2 | 12.4~27.0 (n=3) | 35~253 | 1.3~1.9 | 60.9~85.9/ 49.3~73.5 | 1.1~6.0/ 1.4~5.7/ 1.6~3.9 | 0.5~0.9/ 0.9~5.4 | 79.9~97.6 (E/T=5, 48 hrs) |
|  | 7.7 | Modulator 1 + Modulator 2 + iCasp9-CAR | 11.9, 14.7 (n=2) | 25, 175 | 1.1, 2.3 | 88.5, 94.7/ 85.6, 87.3 | 0.7, 2.6/ 0.8, 3.5/ 1.1, 3.4 | 0, 0.9/ 1.6, 3.7 | 79.9, 85.9 (E/T=5, 48 hrs) |
| GF-CART04 Multiple Solid Tumors | 6.1 | iCasp9-CAR + Modulator 1 | 19.2 (n=1) | 38 | 2.5 | 73.7/ 56.8 | - / 31.1/ 14 | - / 11.3 | 60.2 (E/T=1, 48 hrs) |
|  | 5.1 | iCasp9-CAR + Modulator 2 | 30.7 (n=1) | 139 | 1.9 | 78.8/ 62.6 | - / 13.3/ 3.7 | - / 1.1 | 66.0 (E/T=1, 48 hrs) |

Importantly, we observed no obvious effect of transgene size on the percentage of T_SCM_ cells, which remains the major population in both CD4^+^ or CD8^+^T cell subtypes. Furthermore, in a ten-day culture, using the G-Rex setup with *iCellar^™^* supplementation, the production of clinical scale (up to 10E9) CAR^+^ T_N_/T_SCM_ cells in one liter of culture can still be achieved (data not shown). Together, these results demonstrate that irrespective of gene identity and size, performance of *qCART*^™^-produced T cells is still within satisfactory range. It also lends support to the feasibility of producing multiplexed CAR-T cells for clinical application. It should be noted that the observed downregulation of CAR expression may be due to multiple factors and further experimentation is needed to determine the exact mechanism. Nevertheless, it is important to point out that using the G-Rex setup, we do not observe as dramatic a drop in expression as that seen in virus-transduced CAR-T cells when its payload limit has been reached.

### *qCART^™^* system addresses unmet needs of conventional CAR-T therapies

As evidenced above, *qCART^™^* overcomes major hurdles faced by current virus-based and virus-free genetic engineering and cell production systems. A schematic depiction of *qCART^™^* summarizing and highlighting the roles of each platform is shown in Figure 6.

**Figure 6.**
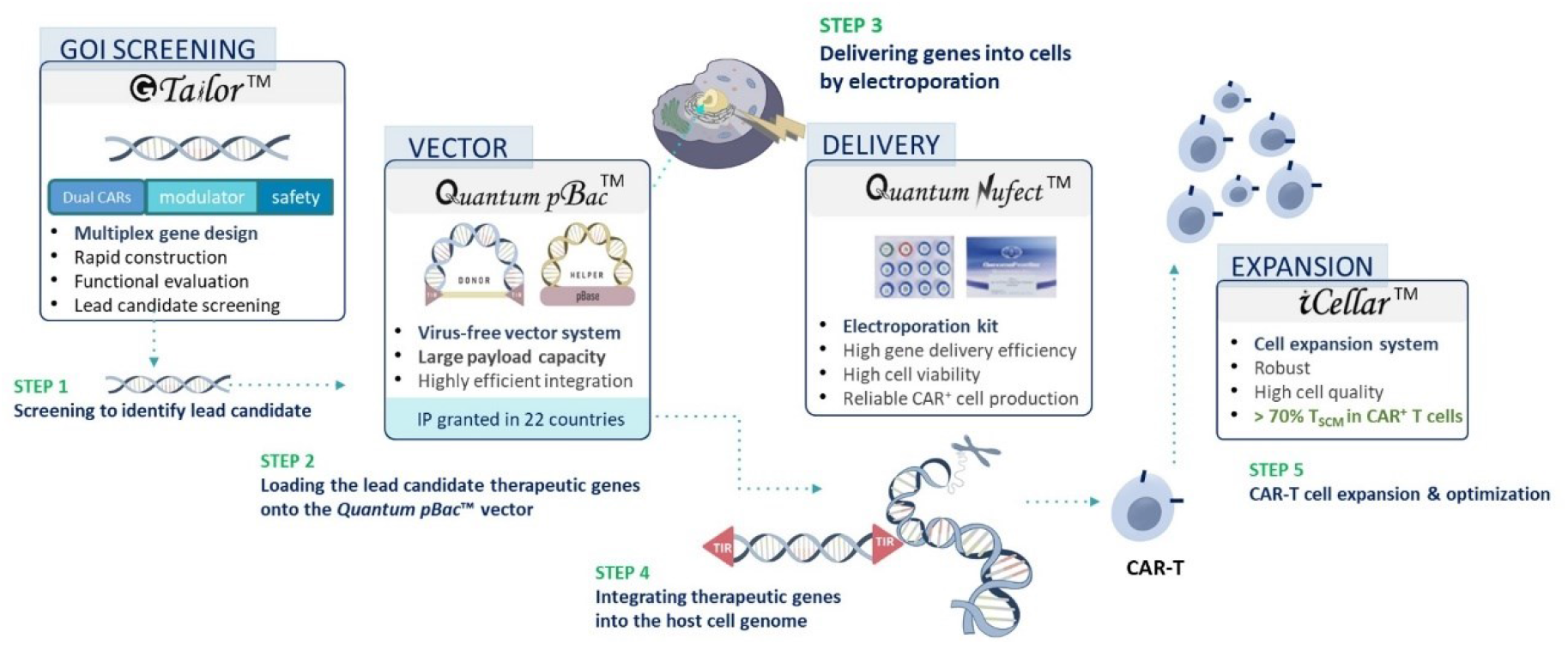
The *qCART^™^* system. A schematic depiction showing the components of the *qCART^™^* system.

## Discussion

In this study, we have shown that the *qCART^™^* system (Figure 6) effectively and robustly produces CAR-T cells having qualities and in quantities that are desirable for clinical application. We have shown that *GTailor^™^* is an efficient multiplex gene design and construct screening platform for designing CAR-T cells with (i) the ability to recognize multiple targets; (ii) modulators for enhancing CAR-T trafficking, infiltration and/or TME resistance; and (iii) a safety switch to terminate the treatment as needed. We also demonstrated that the *qNF* platform is an electroporation-based buffer for introducing therapeutic genes into T cells while preserving cell viability. We previously demonstrated that the *qPB* platform is a virus-free transposon vector system that: (i) possesses a large payload capacity, (ii) is highly efficient in mediating genome integration, and (iii) has high preference for T_SCM_ transposition.^24^ Finally, we showed that *iCellar^™^* is a cell expansion platform for producing clinical-scaled CAR-T cells with (i) high percentage of CAR^+^T_SCM_ cells and (ii) enhanced fitness. In sum, we demonstrated that *qCAR-T^™^* is a streamlined, economic, and robust cell engineering system that expedites the development and manufacturing of virus-free multiplex CAR-T therapy.

To date, the success achieved by CAR-T therapy in B cell lymphomas has not been replicated in solid tumors.^34–36^ The lack of cancer-specific antigens and the immunosuppressive tumor microenvironment are major challenges blocking success in solid tumor. One strategy to overcome these challenges is to engineer T cells to simultaneously express multiple therapeutic genes. This can be achieved using a technology capable of performing one-step multiplex gene integration via a single cargo. Given the complexity of solid tumors and the distinct attributes for each cancer type, a library of CAR-T cells encompassing a large repertoire of GOI designs should be screened to identify the most suitable CAR design. Furthermore, a system for rapid assessment of CAR-T cells’ targeting specificity and On-Target Off-Tumor toxicities is also needed. Lastly, high-throughput cytotoxicity assays to identify the most efficacious CAR-T cell candidates is crucial. *GTailor^™^*, the R&D component of the *qCART^™^* system, addresses all of the aforementioned needs. It streamlines the process for identifying the most optimal multiplex gene design that can be tailored to combat a variety of cancer types.

Consistency during production and in products is the most desirable manufacturing attribute. By applying *qCART*^™^ in small-scale CAR-T cell production, high quality CAR-T products with low batch-to-batch and inter-donor variations are routinely obtained. *qCART*^™^ is able to achieve consistency due to two reasons. The first includes the highly effective transfection, chromosomal transposition, as well as cell expansion during CAR-T cell production process. The second is that the CAR-T product possesses all of the desired features, including reduced safety risks, identity, purity, and potency. The CD19/CD20 tandem CAR-T with iCasp9 is an exemplary product which possesses all of these desired features.^26^ Possible mechanisms that contribute to the desired high CAR-T_SCM_, balanced CD8:CD4 ratio, and reduced safety concerns are discussed below.

A transfection reagent capable of achieving high-efficiency gene delivery with minimal toxicity is imperative for producing reliable gene-modified cell products. We show in this study that, compared to HTCN, nucleofection using *qNF* is a gentler combination that resulted in higher percentages of viable T cells having enhanced expansion capacity (Figures 2 and 3), and higher numbers of CAR-T cells produced (Figure 3). In addition to the known toxic effects of excess DNA^37^ which may contribute to the lower viability of HTCN-transfected CAR-T cells, the observed discrepancy in CAR-T cell production between *qNF- and* HTCN-transfected CAR-T cells may be caused by differential levels of CAR expression. Previous studies have found that high CAR expression on the T cell surface can result in spontaneous antigen-independent clustering of CAR and produce tonic signaling that result in impaired expansion and apoptosis/exhaustion of the cells.^38,39^ In line with this finding, we found that the majority of live HTCN-transfected T cells expressed the transgene at markedly higher (MFI) levels compared with those of *qNF*-transfected T cells (10E2-10E4 and 10E1-10E3, respectively; Figure 2A).^37^ The higher level of CAR expression in HTCN-transfected T cells may have caused greater cellular toxicity, reduced cell survival and subsequent lower CAR-T expansion.^32^ Conversely, the lower CAR expression levels in *qNF*-transfected cells may result in: (1) greater CAR-T cell survival due to reduced toxicity of either DNA or tonic signaling and/or electroporation-induced death and (2) greater CAR^+^ T cell expansion due to lower cell damage after electroporation. The lower transfection rate by *qNF* as compared to HTCN is rescued by *qPB’s* superior gene integration efficiency. Thus, *qNF* and *qPB* platforms cooperate to enrich for CAR^+^ T cells.

Risks associated with random gene integration represent a major safety concern in gene therapy. A recent clinical report demonstrated that 2 of 10 patients treated with *piggyBac-modified* CD19 CAR-T therapy developed CAR-T-cell lymphoma.^40^ A detailed mechanistic study on the case revealed a high number of *piggyBac* integrants, but none were inserted into or near typical oncogenes.^41^ Nevertheless, this study highlights the importance of keeping a low copy number of integrants per genome in gene therapy products. Of note, the study found transgene integration into (introns of) the *BACH2* gene in both malignancies, which we also found in CAR-T cells derived from one of two donors included in the present study (Table S3). However, the study went on to conclude that *BACH2* integration is unlikely to be playing a role in malignant behavior, in part due to *BACH2* also being a recurrent insertion site for retroviruses.^41^

Our previous study has demonstrated that *qPB* is the most effective *piggyBac-based* transposon system for engineering CAR-T cells.^24^ Here, we have further confirmed that *qPB* is a potentially safe vector for producing clinical-grade CAR-T cells given its low copy number of integrant (<5 copies per cell) and similar integration site biodistribution profile compared with other gene engineering and production systems. Furthermore, as compared to *hyPB, qPB* leaves behind significantly lower amount of backbone residues on the genome (107 bp vs. ~600 bp) and lacks the dominant enhancer activity found in backbone residues of *hyPB*, thereby making it a relatively safer *piggyBac* system for CAR-T cell production.^24^

Naïve T cells (T_N_) and T_SCM_ have the capacity to persist in cancer patients, leading to improved clinical outcome.^42^ Hence, enriching these cell types, especially CAR^+^T_SCM_ cells, has become a central focus in the development of next-generation CAR-T therapy.^17–20^ Our studies have demonstrated that almost all *qCART^™^*-generated CAR-T cell products contain >90%T_N_ and T_SCM_ (> 70%T_SCM_ in most cases) in both CD4 and CD8 CAR^+^ T cell populations, regardless of (1) CAR construct design, (2) GOI size, and (3) PBMC source (healthy donors or patients with B-cell malignancies; Figure 4, Table 2).^26^ We believe this desired feature is synergistically achieved by the combined benefits derived from: (1) including 4-1BB rather than CD28 in the CAR construct, (2) using *qNF* instead of HTCN for nucleofection, (3) choosing *qPB* over virus-based or other virus-free (e.g. *hyPB*) vector systems, and (4) using *iCellar^™^, qBT in* particular, for CAR-T cell expansion. Possible mechanisms that may contribute to the high CAR-T_SCM_ enrichment in our CAR-T products include, first, it has been demonstrated that 4-1BB signaling in CAR-T cells promotes T_SCM_ expansion, whereas CD28 favors T_EFF_ expansion.^43^ Second, the magnitude of reduction in cell expansion capacity associated with electroporation-induced damage (Figure 2C) may be amplified in highly proliferating cells such as T_SCM_. Third, extended *ex vivo* expansion of virus-based CAR-T cells often resulted in more differentiated T_EFF_ phenotype with higher expression of exhaustion markers, such as TIM3 and PD1. In contrast, when *ex vivo*-cultured for the same time-frame, *piggyBac-based* CAR-T products consistently have >70% of T_SCM_, which may be further enhanced by using *qPB* rather than *hyPB* as the gene engineering vector system. Finally, current *ex vivo* approaches to expand CAR-T cells to sufficient numbers while maintaining a minimally-differentiated phenotype are hindered by the biological coupling of clonal expansion and effector differentiation. In this study, we demonstrated that *qBT*, a component of *Cellar^™^*, not only promoted robust CAR-T cell expansion but also maintained T cell stemness even in the presence of antigen-expressing aAPC. One possibility is that *qBT* increased the CD4 CAR^+^T_SCM_ population (Figure 4C), which in turn enhanced proliferation of CD8 T_SCM_ population without concomitant differentiation. This may reflect the high (3-9) and balanced (approximately 1)^26^ CD8:CD4 ratios observed in healthy donors and patients, respectively.

Since its first clinical study publication in 2016,^44^ the so called “off-the-shelf” allogeneic CAR-T therapy has been thought of as an inevitable replacement of autologous CAR-T products. However, ample clinical data have concluded that allogeneic CAR-T products are less potent and limited in CAR-T cell persistence as compared to their autologous counterpart. In October 2021, the FDA placed a clinical hold on all Allogene Therapeutics’ AlloCAR-T clinical trials after a chromosomal abnormality was found in a patient who received the anti-CD19 allogeneic CAR-T therapy. The incident further raised concerns regarding all allogeneic CAR-T therapies. Induced pluripotent stem cells (iPSC)-derived CAR-T cells may be a promising alternative for allogeneic CAR-T therapy. However, such approaches are at an early stage of development and require further pre-clinical and clinical research. Furthermore, the heavy mutation burden of iPSCs poses safety concerns regarding its clinical applications. Thus, autologous CAR-T therapy is likely to remain the mainstay of CAR-T based treatment, at least in the short to medium term. In this regard, we have demonstrated that *qCART^™^* addressed most if not all of the current challenges of autologous CAR-T therapy. Thus, we expect this streamlined, robust, and virus-free cell engineering system to unlock the full potential of CAR-T therapy for treating diseases.

## Materials and Methods

### Human T cell samples from healthy donors

Blood samples from adult healthy donors were obtained from Chang Gung Memorial Hospital (Linkou, Taiwan), the acquisition of these samples was approved by the Institution Review Board (IRB No. 201900578A3) at Chang Gung Medical Foundation.

### *GTailor^™^* identification of lead candidates

A proprietary collection of primary cells and (engineered) cell lines were used for *in vitro* evaluation of ON-Target anti-tumor activities. Stable cell lines engineered to express a reporter gene (luciferase and/or GFP or tdTomato) were generated using either a *piggyBac* or a lentivirus vector system. A proprietary collection of parental plasmids of various configurations were used to generate a library of constructs, and the indicated gene(s) of interest (GOI) of various identities and sizes were cloned into the construct’s multiple cloning site. *qPB* parental donor plasmids carrying the indicated GOI were used to construct minicircle DNA using methodology as previously described.^45^ CAR-T cells produced by nucleofection of activated T cells with these minicircle donor constructs in combination with a *Quantum PBase* helper plasmid (i.e. *qPB*) were evaluated for their performance as well as their *in vitro* cytotoxic activity against the collection of cells. Once a lead candidate is identified, its *in vivo* antitumor activity is also evaluated in a mouse xenograft model.

### Mapping of *Quantum pBac^™^* genome integration sites (IS)

Genomic DNA of CAR-T products was extracted using DNeasy Blood and Tissue Kit (QIAGEN, Germantown, MD) and randomly fragmented (<500 bp fragments) using a Bioruptor^®^ Pico sonication device (Diagenode, Denville, NJ). The fragments were ligated to adapters, purified using AMPure XP beads (Beckman Coulter, Indianapolis, IN), and amplified by PCR. The amplified products with an end sequence of the CAR gene were further amplified by PCR. The final PCR products were subjected to sequencing using NovaSeq system (Illumina, San Diego, CA).

Following sequencing, adapter and molecular barcode sequences were removed from the raw reads sequencing data using the Agilent Genomics NextGen Toolkit (AgeNT) software module (Agilent, Santa Clara, CA). The reads were then aligned to the hg38 human genome using Burrows–Wheeler aligner (version 0.7.17-r1188)^46^ and the mapping results sorted using SAMtools (version 1.9).^47^ IS were manually identified using Genome Rearrangement Identification Software Suite (GRIDSS)^48^ and BEDTools (version 2.30.0).^48^ For analysis of IS in proximity to cancer-associated genes, reads data was compared with cancer gene database downloaded from Cancer Hotspots (https://www.cancerhotspots.org/). The next generation sequencing raw data set is shown in Supplementary Table S7.

### Generation and expansion of CAR-T cells

Peripheral blood mononuclear cells (PBMCs) were isolated from blood samples of healthy donors utilizing Ficoll-Hypaque gradient separation. CD3^+^ T cells were isolated from PBMCs using EasySep^™^ Human T Cell Isolation Kit (StemCell Technologies, Taiwan) according to the manufacturer’s instructions. T cells were activated by co-incubation with Dynabeads^™^ (ThermoFisher Scientific, Hillsboro, OR) for two days at a beads-to-cells ratio of 3:1. Following the removal of Dynabeads^™^, activated T cells were subjected to nucleofection with a minicircle CAR donor construct and a *Quantum PBase^™^* plasmid (i.e. *qPB*) using a Nucleofector^™^ 2b or 4D Device (Lonza, Morrisville, NC) in combination with either the Amaxa^®^ Human T Cell Nucleofector^®^ Kit (Lonza) or the *qNF* Kit (GenomeFrontier Therapeutics Inc., Taiwan), as according to the respective manufacturer’s instructions. Nucleofected cells were cultured for 10 or 12 days in OpTmizer medium (Thermo Fisher) supplemented with *qBT* (GenomeFrontier Therapeutics, Taiwan)(and aAPC) in G-Rex 24- or 6M-well plates (Wilson Wolf; “G-Rex setup) or conventional plates (“Perfusion setup”), and then harvested for analysis/further experiments. In experiments assessing the effect of cell viability on expansion capacity, T cells were nucleofected using different program settings (Nucleofector^™^ 2b) and cultured for four days in OpTmizer medium (Thermo Fisher Scientific, Hillsboro, OR) supplemented with 50 IU of IL-2 (PeproTech, Cranbury, NJ) and 10% FBS (Caisson Technologies, Seattle, WA). In experiments involving the addition of aAPC, γ-irradiated aAPC were added on Day 3 to T cells at a 1:1 aAPC:T cell ratio. T cells with or without addition of aAPC were cultured in OpTmizer medium, supplemented with 50 IU of IL-2 and 10% FBS or *qBT*(GenomeFrontier Therapeutics Inc., Taiwan).

### Evaluation of CAR-T cell performance

Unless otherwise specified, flow cytometry analysis performed on a SA3800 Spectral Analyzer (Sony Biotechnology, San Jose, CA) was used to determine the following: CAR expression on T cells was determined by staining with F(ab’)_2_ fragment specific, biotin-conjugated goat anti-mouse antibodies (Jackson ImmunoResearch Laboratories, West Grove, PA), and R-phycoerythrin (PE)-conjugated streptavidin (Jackson ImmunoResearch Laboratories, West Grove, PA). Cells were also stained with one or more of the following antibodies: CD3-Pacific Blue, CD4-Alexa Flour 532 (ThermoFisher Scientific, Hillsboro, OR), CD8-PE-Cy7, CD45RA-BV421, CD62L-PE-Cy5, and CD95-BV711 (Biolegend, San Diego, CA). For determination of live cells, cells were incubated with propidium iodide (PI, Thermo Fisher Scientific, Hillsboro, OR) and/or Acridine orange (AO, Nexcelom, Lawrence, MA). T cell subsets were determined based on CD45RA, CD62L and CD95 staining: T_N_ (CD45RA^+^CD62L^+^CD95^-^), T_SCM_(CD45RA^+^CD62L^+^CD95^+^), T_CM_ (CD45RA^-^CD62L^+^), T_EM_ (CD45RA^-^CD62L^-^), and T_EFF_ (CD45RA^+^CD62L^-^). Histograms and dot-plots were generated using GraphPad Prism software (GraphPad Software, San Diego, CA). Total day 1 live cells were determined using Celigo image cytometry (Nexcelom, Lawrence, MA) and represent the number of AO^+^, PI^-^ cells. Viability index was calculated using the formula: (AO^+^, PI^-^ live cells on day 1/total number of electroporated cells) x 100. Number of CAR^+^ live cells was calculated according to the formula: Total day 1 live cells x percentage of CAR^+^ of live cells.

### *In vivo* cytokine release assay

Mouse serum samples were collected on Days 3, 15, 29 and 43 following T cell injection. Serum levels of interferon gamma (IFN-γ) was measured by performing enzyme-linked immunosorbent assay (Thermo Fisher Scientific, Hillsboro, OR) according to the manufacturer’s instructions.

### *In vitro* cytotoxicity assay

5×10^3^ cells per well of Raji-GFP/Luc target cells were seeded in 96-well culture plates (Corning, Glendale, AZ) and CAR-T cells were added at an E:T ratio of 1:1. CAR-T cell-mediated cytotoxicity on Raji-GFP/Luc target cells was then assessed by using Celigo image cytometry (Nexcelom, Lawrence, MA) as reported^49^ to determine the number of live Raji-GFP/Luc cells at 0, 24, 48, 72 and 96 hours after co-culturing. Cell aggregates were separated by pipetting before Celigo imaging. The percentage of specific lysis for each sample was calculated using the formula: [1-(live fluorescent cell count in the presence of Raji-GFP/Luc cells and CAR-T cells / live fluorescent cell count in the presence of Raji-GFP/Luc cells only)] x 100.

### Mouse xenograft model

*In vivo* studies using mouse xenograft model were conducted at the Development Center for Biotechnology, Taiwan, using animal protocols approved by the Taiwan Mouse Clinic IACUC (2020-R501-035). Briefly, eight-week-old female ASID (NOD.Cg-Prkdc^scid^Il2rg^tm1Wjl^/YckNarl) mice (National Laboratory

Animal Center, Taiwan) were intravenously (i.v.) injected with 1.5×10^5^ Raji-GFP/Luc tumor cells. One week after Raji-GFP/Luc tumor cell injection, mice were injected with 3×10^6^ CAR-T cells or control Pan-T cells (non-transfected T cells). Luminescence signals from Raji-GFP/Luc tumor cells were monitored using the Xenogen-IVIS Imaging System (Caliper Life Sciences, Hopkinton, MA).

### Genomic DNA extraction and quantitative PCR (qPCR)

Genomic DNA from mouse blood was extracted using DNeasy Blood & Tissue Kit (Qiagen, Germantown, MD) following the manufacturer’s instructions. 7500 fast real-time PCR system (Applied Biosystems, Foster City, CA) was used to carry out quantitative PCR analysis. Amplification of CAR gene (CAR-T cells) and Luc gene (Raji-GFP/Luc cells) in mouse blood samples was carried out using the CAR forward (5’-ACGTCGTACTCTTCCCGTCT-3’) and reverse (5’-GATCACCCTGTAC-TGCAACCA-3’) primers and the luciferase forward (5’-GGACTTGGACACCGGTAAGA-3’) & reverse (5’-GGTCCACGATGAAGAAGTGC-3’) primers, respectively. The amount of CAR and Luc genes, expressed as gene copy/ng of DNA, were calculated utilizing the standard curve method as previously described.^50^

### Statistical analysis

Statistical analyses of differences between two groups and among three or more groups were carried out using the Student’s t-test (two-tailed) and the one-way ANOVA with Tukey’s multiple comparison test, respectively. The analyses were performed using Prism 7.0 (GraphPad Software, San Diego, CA), and statistical significance was reported as * *p*<0.05, ** *p*<0.01, and *** *p*<0.001.

## Acknowledgments

The authors thank Ms. Yi-Shan Yu and Ms. Lu-Chun Chen for their assistance throughout the IRB preparation and approval process. The authors also thank Dr. Pei-Yi Tsai for assistance with the animal experiments. This study is funded by GenomeFrontier Therapeutics, Inc.

## Author Contributions

S.C.-Y.W. designed research. Y.-C.C., W.-K.H., Y.-W.H., J.-C.T., Y.-H.K., P.-H.W., P.-N.W., K.-F.C., W.-T.L. performed research. J.C.H., Y.-C.C, W.-K.H., K.-L.K.W., and S.C.-Y.W. analyzed data. J.C.H., P.S.C. and S.C.-Y.W. wrote the paper.

