## Supplementary Materials for "*Quantum CART* (*qCART*), a *piggyBac-based* system for development and production of virus-free multiplex CAR-T cell therapy"

**This supplementary information file contains:**

(1) Supplementary Figure Legends; and

(2) Figures S1-S5, and Tables S1-S7

**Supplementary Figure Legend**

**Supplementary Figure S1. Screening of phage library and binding specificity confirmation to select for candidate CAR binding domain.** Flow chart of selection procedure utilized to screen for candidate CAR binding domains for pipeline production.

**Supplementary Figure S2. Higher rate of increase of CAR$^+$ T cells obtained from human primary T cells electroporated using the *Quantum Nufect*™ (*qNF*) buffer system compared with those electroporated using the Human T Cell Nucleofector™ (HTCN) solution.** (A) Human peripheral blood T cells electroporated with *Quantum pBac*™ expressing CAR in either *qNF* buffer or HTCN solution were analyzed one day and 14 days after electroporation for percentages of CAR$^+$ cells. (B) Rate of changes in % CAR$^+$ cells. Horizontal lines (B) represent median percentage of CAR$^+$ cells. Each panel of results shown in (A) is data of one individual donor. Rate of changes in % $CAR^+$ cells is calculated using the formula: (% $CAR^+$ cells on day 14 - % $CAR^+$ cells on day 1) / total number of days in culture. N = 7 donors.

**Supplementary Figure S3. Profiling of mice treated with CAR-T cells produced using the *qCART*™ system.** (A) Plasma concentration of IFN-$\gamma$, (B) the copy number of CAR per ng of non-RBCs in blood, and (C) luciferase activity as a reporter for Raji cells in the blood of mice on day 85, were determined. Results shown are from four to eight mice/group. Pan-T cells were used as a control. *Quantum Booster*™ (*qBT*) was present in all cell culture conditions. Dotted line (B) represents limit of detection. * $p < 0.05$.

**Supplementary Figure S4. Determination of T cell subsets in CAR-T cells produced using the *qCART*™ system.** Human peripheral blood mononuclear cells (PBMC) with or without pre-incubation with aAPC and electroporated with *Quantum pBac*™ expressing CAR were analyzed for their T cell subset distribution profile (percentages of $T_N$/$T_{SCM}$, $T_{CM}$, $T_{EM}$ and $T_{EFF}$ cell subsets in $CAR^+CD4^+$ (left panel) or $CAR^+CD8^+$ (right panel) cells). Dot plot and histogram data shown are from one staining experiment of CAR-T cells produced from PBMC of healthy donor A.

**Supplementary Figure S5. Determination of CAR copy numbers in T cells produced using the *qCART*™ system.** Copy numbers of the CAR gene per T cell from T (Donor

1; left panel) and T (Donor 2; right panel) were analyzed using digital PCR. Results shown are from one representative experiment.

**Figure S1**

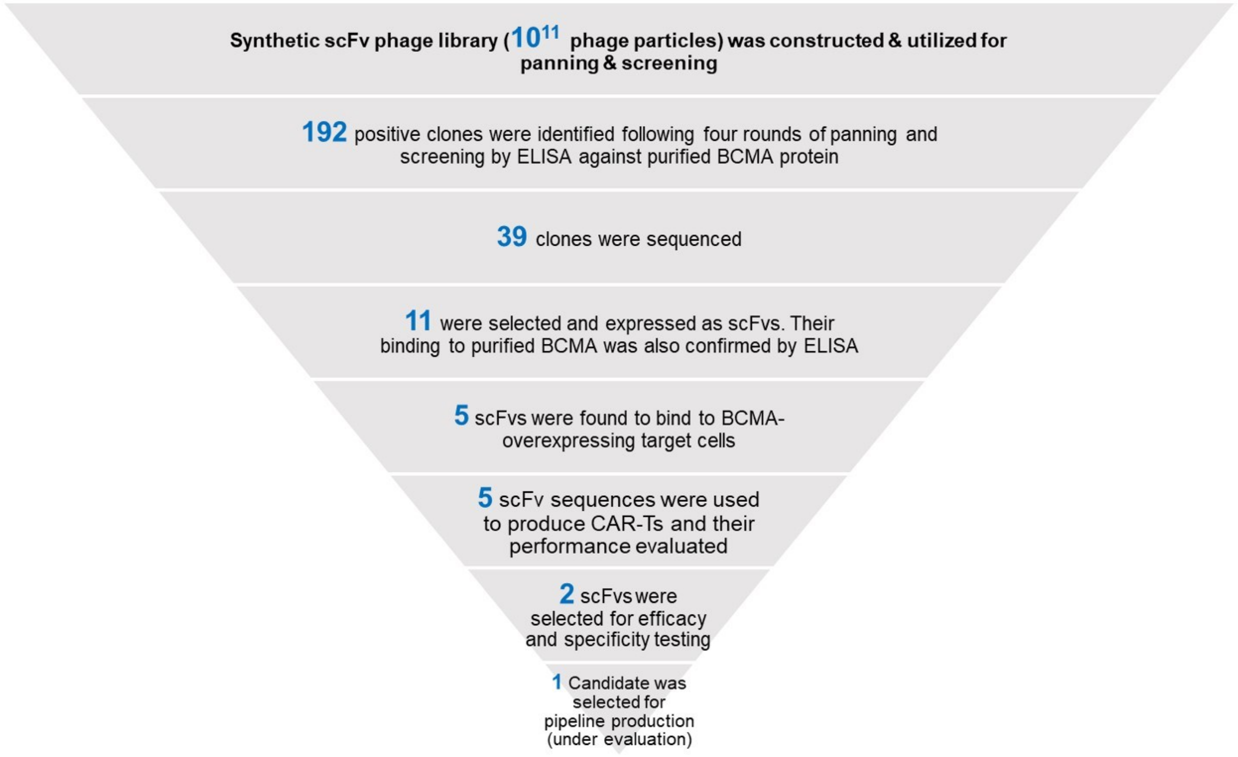
Synthetic scFv phage library (10^11 phage particles) was constructed & utilized for panning & screening
192 positive clones were identified following four rounds of panning and screening by ELISA against purified BCMA protein
39 clones were sequenced
11 were selected and expressed as scFvs. Their binding to purified BCMA was also confirmed by ELISA
5 scFvs were found to bind to BCMA-overexpressing target cells
5 scFv sequences were used to produce CAR-Ts and their performance evaluated
2 scFvs were selected for efficacy and specificity testing
1 Candidate was selected for pipeline production (under evaluation)

### Figure S2

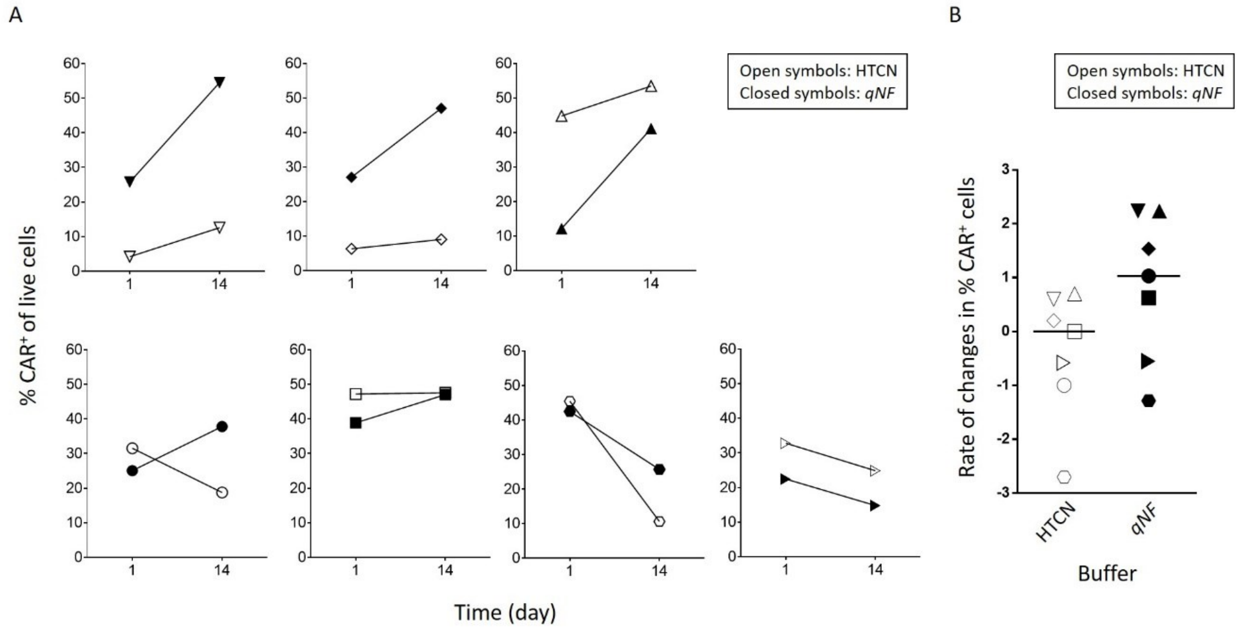
A
B
Open symbols: HTCN
Closed symbols: qNF
Open symbols: HTCN
Closed symbols: qNF
% CAR+ of live cells
Time (day)
Rate of changes in % CAR+ cells
HTCN
qNF
Buffer

### Figure S3

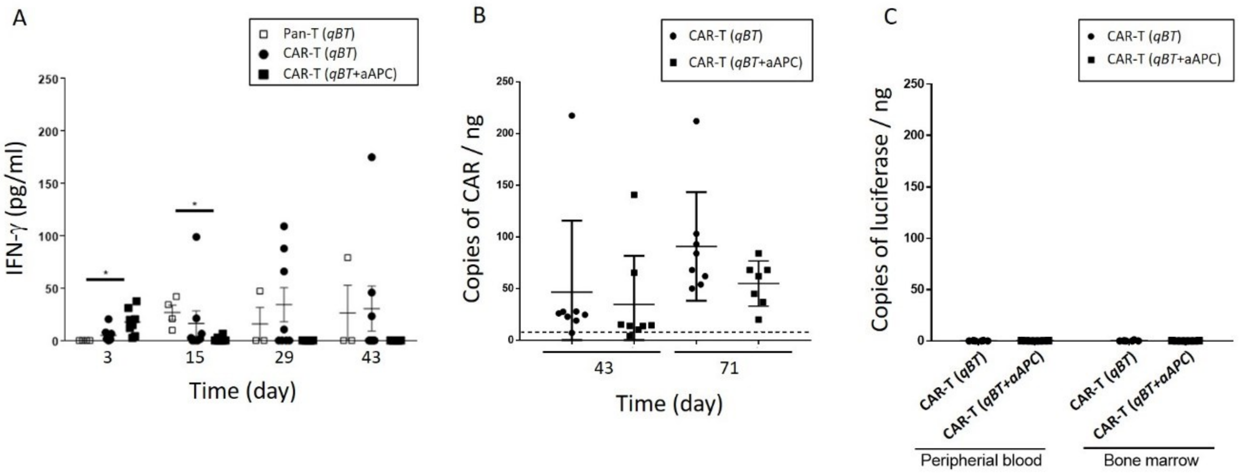
A
Pan-T (qBT)
CAR-T (qBT)
CAR-T (qBT+aAPC)
IFN-γ (pg/ml)
Time (day)
3
15
29
43
B
CAR-T (qBT)
CAR-T (qBT+aAPC)
Copies of CAR / ng
43
71
Time (day)
C
CAR-T (qBT)
CAR-T (qBT+aAPC)
Copies of luciferase / ng
CAR-T (qBT)
CAR-T (qBT+aAPC)
CAR-T (qBT)
CAR-T (qBT+aAPC)
Peripherial blood
Bone marrow

**Figure S4**

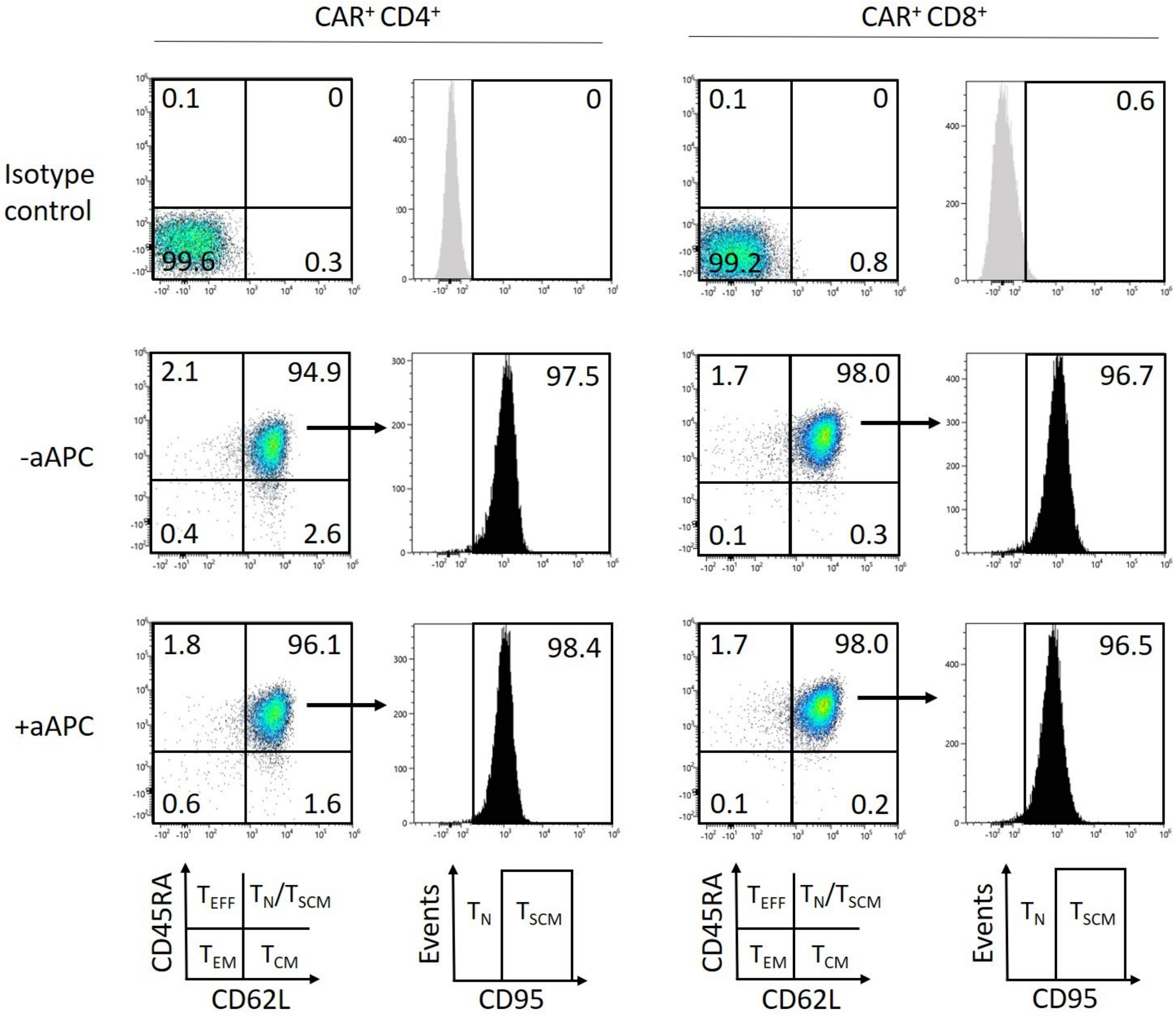
CAR+ CD4+
CAR+ CD8+
Isotype control
-aAPC
+aAPC
0.1
0
99.6
0.3
0
0.1
0
99.2
0.8
0.6
2.1
94.9
0.4
2.6
97.5
1.7
98.0
0.1
0.3
96.7
1.8
96.1
0.6
1.6
98.4
1.7
98.0
0.1
0.2
96.5
CD45RA
CD62L
TEFF
TN/TSCM
TEM
TCM
Events
CD95
TN
TSCM

### Figure S5

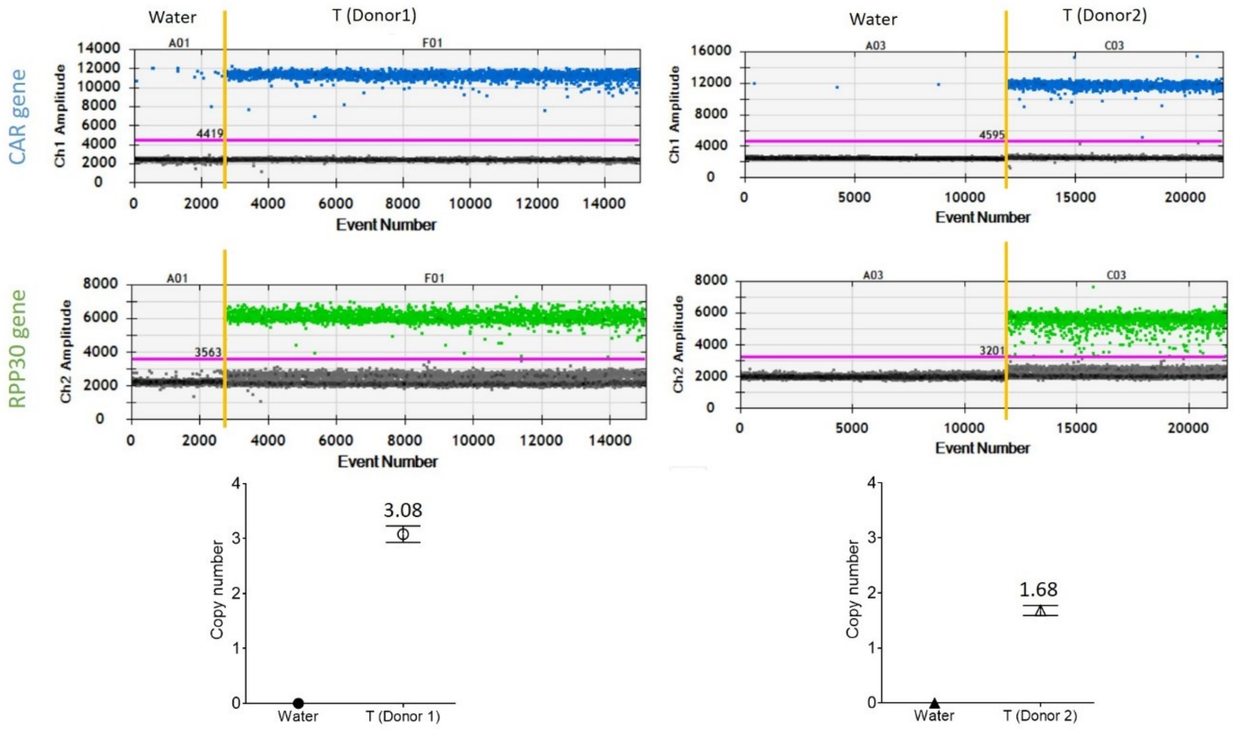

Water
T (Donor1)
CAR gene
Ch1 Amplitude
A01
F01
4419
Event Number
Water
T (Donor2)
A03
C03
4595
RPP30 gene
Ch2 Amplitude
3563
3201
Copy number
3.08
1.68
Water
T (Donor 1)
T (Donor 2)

### Table S1. *In vitro* cytotoxicity of various human target cells and cell lines

| No. | CAR-T | % lysis of GFP$^{+}$ target cells (E:T=1:1, 48 h) | | | | | | | | | | | | | | | | % lysis of CMFDA-labelled target cells (E:T=5:1 48 h) | | | |
|---|---|---|---|---|---|---|---|---|---|---|---|---|---|---|---|---|---|---|---|---|---|
| | | BCMA negative control | BCMA positive control | Multiple myeloma | | Leukemia | | Breast cancer | | | | Pancreatic cancer | Ovarian cancer | Stomach cancer | Lung cancer | Glioblastoma | Normal MSC | Normal cells | | | |
| | | | | A | B | A | B | A | B | C | D | A | A | A | A | A | A | Primary fibroblasts | MSC 1 | MSC 2 | PBMC |
| (%BCMA) | | 0.1 | 90 | 79.5 | 93.4 | 0.8 | 3.3 | 0.2 | 1.3 | 0.2 | 0.1 | 0.4 | 0.1 | 0.4 | 0.1 | 0.9 | 0.4 | 1.7 | 0.2 | 9.3 | 0.6 |
| 1 | aBCMA-2 | +/- | +++ | +++ | +++ | - | +/- | - | - | - | - | +/- | +/- | +/- | - | - | - | - | - | - | - |
| 2 | aBCMA-5 | +/- | +++ | ++ | +++ | +/- | - | +/- | - | - | - | + | - | + | +/- | +/- | - | +/- | - | - | - |

- <0%; +/- 0~10%; + 11~20%; ++ 21~30%; +++ 31~50%; ++++ >51%

### Table S2. Summary of criteria utilized for identification of insertion sites

| T cell donor | Copy number | Raw reads | No. of reads containing CAR and human sequence (>25bp) | No. of reads mapping to integration sites | No. of reads mapping to unique integration sites |
|---|---|---|---|---|---|
| 1 (IRB1) | 3.08 | 8,463,487 | 512,708 | 511,824 | 344 |
| 2 (IRB23) | 1.68 | 5,659,685 | 287,521 | 286,812 | 300 |

**Table S3. Top 10 gene clusters containing IS**

IS data is analyzed and grouped as according to the top 10 protein-coding gene clusters within each of which IS were found. Results are presented as percentages of total IS.

| | T (Donor 1) | | T (Donor 2) | |
|---|---|---|---|---|
| Rank | Gene | Frequency (%) | Gene | Frequency (%) |
| Top 1 | DOCK8 | 1.56% | RAB27A | 1.86% |
| Top 2 | SLC1A5 | 1.10% | BCAS4 | 1.13% |
| Top 3 | BCKDHB | 0.91% | BACH2 | 1.10% |
| Top 4 | ZNF320 | 0.88% | ENOX1 | 1.07% |
| Top 5 | WDR17 | 0.87% | OR7D2 | 0.98% |
| Top 6 | NDUFB3 | 0.80% | AMD1 | 0.96% |
| Top 7 | DCAF7 | 0.79% | LCA5L | 0.96% |
| Top 8 | NDUFAF2 | 0.77% | TNFAIP8 | 0.95% |
| Top 9 | MGAT4A | 0.76% | MELTF | 0.95% |
| Top 10 | FDX1 | 0.74% | PBLD | 0.90% |
| All other mapped IS | | 90.83% | | 89.14% |

**Table S4. IS found within/nearby cancer-associated genes**

IS data is analyzed and grouped as according to the cancer-associated genes within/near each of which IS were found. Results are presented as percentages of total IS.

| Sample ID | IS in proximity to cancer-associated gene* | Chromosome | Frequency (%) | Integration region |
|---|---|---|---|---|
| T (Donor 1) | CBL | chr11 | 0.16% | Intron |
| | BCL2 | chr18 | 0.29% | Intron |
| | SDHA | chr5 | 0.01% | Intron |
| | ATRX | chrX | 0.18% | Intron |
| T (Donor 2) | PIK3CD | chr1 | 0.42% | Intron |
| | PIK3CD | chr1 | 0.24% | Intron |
| | PIK3R2 | chr19 | 0.02% | Upstream of gene (<5kb) |
| | IL7R | chr5 | 0.48% | Intron |
| | CDK6 | chr7 | 0.31% | 3_prime_UTR_variant |
| | RHEB | chr7 | 0.05% | Intron |
| | KMT2C | chr7 | 1.10% | Upstream of gene (<5kb) |

* Clustered integration was not found in or nearby genes (*CCND2, HMGA2, LMO2, MECOM, MN1, PRDM16*) previously reported to be associated with severe adverse events in patients

**Table S5. Performance of CAR-T cells derived from human PBMC1996**

Human peripheral blood mononuclear cells (PBMC) with or without pre-incubation with aAPC and electroporated with *Quantum pBac*™ expressing CAR were analyzed for their performance, including cell expansion fold change, percentage of $CAR^+$ cells, and percentage of $T_{SCM}$ cell subsets in $CD4^+$ or $CD8^+$ cells. Data shown are from four healthy donors (A, B, C, and D, respectively).

| Performance | aAPC | Donor | | | |
|---|---|---|---|---|---|
| | | A | B | C | D |
| Fold change | - | 472.44 | 629.03 | 1035.79 | 688.29 |
| | + | 983.27 | 1714.03 | 1554.9 | 1294.93 |
| % $CAR^+$ | - | 53.91 | 36.83 | 29.87 | 33.97 |
| | + | 66.48 | 67.79 | 59.87 | 54.93 |
| % $CD4^+$ $T_{SCM}$ cells | - | 92.36 | 84.22 | 77.26 | 78.44 |
| | + | 92.66 | 80.94 | 79.41 | 79.48 |
| % $CD8^+$ $T_{SCM}$ cells | - | 93.3 | 90.06 | 80.1 | 79.31 |
| | + | 92.88 | 82.14 | 82.37 | 78.52 |

**Table S6. Evaluation of different transgene sizes on CAR-T cell performance**

Donor T cells were electroporated with *Quantum pBac™* expressing transgenes with sizes ranging from 3.0 kb to 7.6 kb. The percentages of $CAR^+$ T cells, expansion fold and relative distribution of $T_{SCM}$ cell subsets in $CD4^+$ $CAR^+$ and $CD8^+$ $CAR^+$ T cell subtypes are shown. Data shown are from 2 to 11 healthy donors.

| | Multiple Myeloma (MM) | | Hematopoietic B Cell Malignancy | | | |
|---|---|---|---|---|---|---|
| Transgene (size) | Target X (3.0 kb) | Target Y (3.0 kb) | CD20/CD19 (3.7 kb) | CD20/CD19 + iCasp9 (5.2 kb) | CD20/CD19 + modulator A (5.4 kb) | CD20/CD19 + modulator B (7.6 kb) |
| Number of donors tested | 2 | 2 | 11 | 10 | 3 | 3 |
| % $CAR^+$ | 77.3 ~ 83.9 | 80.6 ~ 88.3 | 54.4 ~ 79.0 | 44.3 ~ 67.8 | 42.0 ~ 47.7 | 28.7 ~ 42.7 |
| Expansion fold | 1,507 ~ 1,910 | 1,790 ~ 4,190 | 103 ~ 1,501 | 406 ~ 2,090 | 85 ~ 315 | 75 ~ 190 |
| % $T_{SCM}$<br>in $CD4^+$:<br>In $CD8^+$: | 68.0 ~ 86.8<br>67.2 ~ 77.2 | 74.3 ~ 82.4<br>76.5 ~ 76.5 | 18.2 ~ 65.8<br>45.8 ~ 83.4 | 65.1 ~ 92.7<br>66.7 ~ 92.9 | 46.2 ~ 61.9<br>57.9 ~ 70.6 | N.D.<br>62.5 ~ 74.1 |

**Table S7. Next Generation Sequencing (NGS) raw dataset (please see dataset file separately attached)**
