## Supplementary material for "*Quantum CART* (*qCART*), a *piggyBac-based* system for development and production of virus-free multiplex CAR-T cell therapy": Table S7

| Table S7. Next Generation Sequencing (NGS) raw dataset | | | | | | | | |
|---|---|---|---|---|---|---|---|---|
| ID | #CHROM | CytoBand | Position | Reads count | Gene insertion | Genome region | CpG.name | CpG Distance |
| Donor1_001 | chr1 | q44 | 247,548,450 | 253 | GCSAML | upstream_gene_variant | CpG: 51 | 17251 |
| Donor1_002 | chr1 | q43 | 238036571 | 2855 | | intergenic_variant | CpG: 22 | 252423 |
| Donor1_003 | chr1 | q43 | 237197980 | 1097 | RYR2 | intron_variant | CpG: 169 | 154636 |
| Donor1_004 | chr1 | q42.3 | 234723794 | 732 | | downstream_gene_variant | CpG: 329 | 112516 |
| Donor1_005 | chr1 | q42.2 | 232312617 | 1591 | LOC101929902 | intron_variant&non_coding_transcript_variant | CpG: 173 | 316726 |
| Donor1_006 | chr1 | q42.2 | 231556219 | 279 | TSNAX | intron_variant | CpG: 56 | 27357 |
| Donor1_007 | chr1 | q42.2 | 231565892 | 24 | TSNAX | 3_prime_UTR_variant | CpG: 56 | 37030 |
| Donor1_008 | chr1 | q42.12 | 226779892 | 1133 | RPS27P5 | upstream_gene_variant | CpG: 171 | 41040 |
| Donor1_009 | chr1 | q42.12 | 225780514 | 947 | SRP9 | intron_variant | CpG: 35 | 2482 |
| Donor1_010 | chr1 | q32.3 | 211713088 | 465 | | downstream_gene_variant | CpG: 43 | 37254 |
| Donor1_011 | chr1 | q32.3 | 213047287 | 1349 | RPS6KC1 | upstream_gene_variant | CpG: 27 | 3983 |
| Donor1_012 | chr1 | q32.1 | 200153645 | 701 | NR5A2 | intron_variant | CpG: 45 | 5569 |
| Donor1_013 | chr1 | q31.3 | 198244413 | 1784 | NEK7 | intron_variant | CpG: 21 | 85179 |
| Donor1_014 | chr1 | q31.2 | 193014357 | 1030 | UCHL5 | 3_prime_UTR_variant | CpG: 94 | 44735 |
| Donor1_015 | chr1 | q31.1 | 189118036 | 59 | | intergenic_variant | CpG: 23 | 2437087 |
| Donor1_016 | chr1 | q25.3 | 184041907 | 395 | COLGALT2 | upstream_gene_variant | CpG: 140 | 4275 |
| Donor1_017 | chr1 | q24.2 | 168517920 | 1650 | | intergenic_variant | CpG: 87 | 291319 |
| Donor1_018 | chr1 | q24.2 | 168416520 | 1033 | | intron_variant&non_coding_transcript_variant | CpG: 87 | 189919 |
| Donor1_019 | chr1 | q23.3 | 162034919 | 374 | | upstream_gene_variant | CpG: 100 | 10578 |
| Donor1_020 | chr1 | p36.11 | 24511396 | 2284 | RCAN3 | intron_variant | CpG: 108 | 8008 |
| Donor1_021 | chr1 | p36.11 | 23,606,222 | 1463 | MDS2 | intron_variant&non_coding_transcript_variant | CpG: 66 | 37648 |
| Donor1_022 | chr1 | p35.2 | 32256598 | 1008 | LCK | intron_variant | CpG: 130 | 15058 |
| Donor1_023 | chr1 | p34.3 | 38553302 | 303 | | intergenic_variant | CpG: 42 | 25192 |
| Donor1_024 | chr1 | p34.3 | 39196418 | 768 | MACF1 | intron_variant | CpG: 142 | 90231 |
| Donor1_025 | chr1 | p32.3 | 52551428 | 873 | TUT4 | intron_variant | CpG: 115 | 1159 |
| Donor1_026 | chr1 | p32.3 | 52,765,376 | 827 | ZYG11B | intron_variant | CpG: 46 | 38620 |
| Donor1_027 | chr1 | p31.3 | 63322540 | 1301 | FOXD3 | upstream_gene_variant | CpG: 599 | 0 |
| Donor1_028 | chr1 | p31.3 | 63507250 | 957 | ITGB3BP | intron_variant | CpG: 99 | 86017 |
| Donor1_029 | chr1 | p22.1 | 92473204 | 669 | GFI1 | 3_prime_UTR_variant | CpG: 494 | 7106 |
| Donor1_030 | chr1 | p21.2 | 101136905 | 1700 | LOC107985094 | intron_variant&non_coding_transcript_variant | CpG: 26 | 99325 |
| Donor1_031 | chr1 | p13.3 | 109095642 | 1291 | TMEM167B | 3_prime_UTR_variant | CpG: 74 | 4400 |
| Donor1_032 | chr1 | p13.3 | 110903378 | 407 | CD53 | downstream_gene_variant | CpG: 87 | 59882 |
| Donor1_033 | chr1 | p13.2 | 114512497 | 3101 | TRIM33 | upstream_gene_variant | CpG: 129 | 854 |
| Donor1_034 | chr1 | p13.2 | 114512512 | 3101 | TRIM33 | upstream_gene_variant | CpG: 129 | 869 |
| Donor1_035 | chr1 | p13.2 | 114512513 | 3101 | TRIM33 | upstream_gene_variant | CpG: 129 | 870 |
| Donor1_036 | chr1 | p12 | 120176628 | 386 | SEC22B | upstream_gene_variant | CpG: 32 | 94 |
| Donor1_037 | chr1 | p12 | 120176629 | 386 | SEC22B | upstream_gene_variant | CpG: 32 | 95 |
| Donor1_038 | chr10 | q24.32 | 103078196 | 1119 | CNNM2 | 3_prime_UTR_variant | CpG: 117 | 114290 |
| Donor1_039 | chr10 | q24.1 | 95859101 | 992 | | upstream_gene_variant | CpG: 69 | 48386 |
| Donor1_040 | chr10 | q23.32 | 91241466 | 2544 | PCGF5 | intron_variant | CpG: 57 | 20258 |
| Donor1_041 | chr10 | q22.2 | 74157292 | 76 | ADK | intron_variant | CpG: 50 | 5570 |
| Donor1_042 | chr10 | p15.1 | 5842500 | 156 | GDI2 | upstream_gene_variant | CpG: 23 | 1828 |
| Donor1_043 | chr10 | p14 | 7787453 | 2185 | KIN | intron_variant | CpG: 77 | 173 |
| Donor1_044 | chr10 | p13 | 16441414 | 661 | PTER | intron_variant | CpG: 40 | 4110 |
| Donor1_045 | chr10 | p12.31 | 20972852 | 778 | NEBL | intron_variant | CpG: 149 | 200348 |
| Donor1_046 | chr11 | q24.3 | 129955142 | 78 | PRDM10 | intron_variant | CpG: 119 | 47086 |
| Donor1_047 | chr11 | q23.3 | 117976687 | 206 | | intergenic_variant | CpG: 62 | 9237 |
| Donor1_048 | chr11 | q23.3 | 119263569 | 650 | CBL | intron_variant | CpG: 19 | 47118 |
| Donor1_049 | chr11 | q23.3 | 119178865 | 127 | NLRX1 | intron_variant | CpG: 53 | 9630 |
| Donor1_050 | chr11 | q23.1 | 111971264 | 458 | DIXDC1 | intron_variant | CpG: 45 | 5940 |
| Donor1_051 | chr11 | q22.3 | 110433383 | 3103 | FDX1 | intron_variant | CpG: 91 | 2773 |
| Donor1_052 | chr11 | q22.3 | 104689571 | 1114 | LOC105369466 | intron_variant&non_coding_transcript_variant | CpG: 54 | 525241 |
| Donor1_053 | chr11 | q21 | 93540590 | 2512 | SMCO4 | intron_variant | CpG: 116 | 2059 |
| Donor1_054 | chr11 | q21 | 95251530 | 2466 | | intergenic_variant | CpG: 23 | 19106 |
| Donor1_055 | chr11 | q14.2 | 86165002 | 1545 | | regulatory_region_variant | CpG: 18 | 79342 |
| Donor1_056 | chr11 | q14.2 | 87240948 | 573 | TMEM135 | intron_variant | CpG: 55 | 202897 |
| Donor1_057 | chr11 | q13.5 | 76741787 | 1600 | | intergenic_variant | CpG: 149 | 41345 |
| Donor1_058 | chr11 | q13.4 | 71786302 | 1036 | LOC105379409 | non_coding_transcript_exon_variant | CpG: 121 | 224 |
| Donor1_059 | chr11 | q13.1 | 66064531 | 3025 | SF3B2 | intron_variant | CpG: 139 | 5253 |
| Donor1_060 | chr11 | q12.3 | 62728755 | 1586 | TTC9C | 5_prime_UTR_variant | CpG: 142 | 1100 |
| Donor1_061 | chr11 | p15.4 | 10406609 | 1290 | CAND1.11 | intron_variant&non_coding_transcript_variant | CpG: 80 | 43845 |
| Donor1_062 | chr11 | p15.2 | 14684363 | 2113 | PDE3B | intron_variant | CpG: 24 | 39235 |
| Donor1_063 | chr11 | p15.2 | 15059443 | 734 | CALCB | intron_variant | CpG: 69 | 13969 |
| Donor1_064 | chr11 | p15.2 | 16511697 | 688 | SOX6 | intron_variant | CpG: 101 | 92810 |
| Donor1_065 | chr11 | p14.3 | 22314008 | 288 | | intron_variant&non_coding_transcript_variant | CpG: 38 | 27309 |
| Donor1_066 | chr11 | p14.1 | 28021544 | 891 | KIF18A | intron_variant | CpG: 25 | 88773 |

| | | | | | | | | |
|---|---|---|---|---|---|---|---|---|
| Donor1_067 | chr11 | p13 | 32337880 | 1786 | THEM7P | intron_variant&non_coding_transcript_variant | CpG: 63 | 3942 |
| Donor1_068 | chr12 | q24.33 | 133177158 | 873 | ZNF268 | upstream_gene_variant | CpG: 24 | 4203 |
| Donor1_069 | chr12 | q24.31 | 122393528 | 1761 | CLIP1 | intron_variant | CpG: 140 | 28330 |
| Donor1_070 | chr12 | q24.31 | 120529625 | 1432 | COQ5 | upstream_gene_variant | CpG: 27 | 390 |
| Donor1_071 | chr12 | q24.31 | 121724519 | 198 | TMEM120B | intron_variant | CpG: 49 | 11334 |
| Donor1_072 | chr12 | q24.23 | 120116024 | 1427 | RAB35 | intron_variant | CpG: 121 | 335 |
| Donor1_073 | chr12 | q24.23 | 118415125 | 206 | SUDS3 | 3_prime_UTR_variant | CpG: 99 | 38091 |
| Donor1_074 | chr12 | q24.13 | 112227796 | 329 | HECTD4 | intron_variant | CpG: 48 | 42785 |
| Donor1_075 | chr12 | q23.3 | 108584710 | 406 | | intergenic_variant | CpG: 16 | 7139 |
| Donor1_076 | chr12 | q23.1 | 96212268 | 748 | ELK3 | intron_variant | CpG: 133 | 16901 |
| Donor1_077 | chr12 | q23.1 | 96196871 | 413 | ELK3 | intron_variant | CpG: 133 | 1504 |
| Donor1_078 | chr12 | q22 | 95553894 | 1549 | USP44 | upstream_gene_variant | CpG: 137 | 4691 |
| Donor1_079 | chr12 | q21.33 | 89155177 | 1550 | LINC02458 | intron_variant&non_coding_transcript_variant | CpG: 248 | 196215 |
| Donor1_080 | chr12 | q21.33 | 91948057 | 550 | LOC105369901 | intron_variant&non_coding_transcript_variant | CpG: 128 | 196878 |
| Donor1_081 | chr12 | q21.2 | 75975845 | 927 | | intron_variant&non_coding_transcript_variant | CpG: 142 | 55219 |
| Donor1_082 | chr12 | q14.3 | 66599239 | 814 | GRIP1 | intron_variant | CpG: 65 | 363615 |
| Donor1_083 | chr12 | q14.1 | 62259516 | 1880 | USP15 | upstream_gene_variant | CpG: 74 | 569 |
| Donor1_084 | chr12 | q14.1 | 62259562 | 1880 | USP15 | upstream_gene_variant | CpG: 74 | 523 |
| Donor1_085 | chr12 | q13.2 | 55879438 | 2461 | | intergenic_variant | CpG: 24 | 47518 |
| Donor1_086 | chr12 | q13.13 | 53378882 | 3767 | SP1 | upstream_gene_variant | CpG: 52 | 1454 |
| Donor1_087 | chr12 | q13.13 | 54025214 | 1495 | | intron_variant | CpG: 31 | 4430 |
| Donor1_088 | chr12 | q12 | 42092712 | 307 | GXYLT1 | intron_variant | CpG: 92 | 51612 |
| Donor1_089 | chr12 | p13.33 | 539534 | 727 | B4GALNT3 | intron_variant | CpG: 35 | 30260 |
| Donor1_090 | chr12 | p13.33 | 774642 | 175 | WNK1 | intron_variant | CpG: 197 | 20395 |
| Donor1_091 | chr12 | p13.31 | 6424481 | 1170 | | intergenic_variant | CpG: 55 | 40125 |
| Donor1_092 | chr12 | p13.31 | 6574765 | 271 | | intron_variant&NMD_transcript_variant | CpG: 70 | 6176 |
| Donor1_093 | chr12 | p13.31 | 6446883 | 1083 | CD27 | intron_variant | CpG: 76 | 23507 |
| Donor1_094 | chr12 | p13.2 | 10405405 | 1012 | KLRC4 | downstream_gene_variant | CpG: 55 | 191939 |
| Donor1_095 | chr12 | p13.1 | 12768859 | 1255 | STX8P1 | downstream_gene_variant | CpG: 71 | 17921 |
| Donor1_096 | chr12 | p12.3 | 19422251 | 3328 | | upstream_gene_variant | CpG: 147 | 16999 |
| Donor1_097 | chr12 | p11.21 | 31744702 | 1174 | LOC105369724 | downstream_gene_variant | CpG: 51 | 15175 |
| Donor1_098 | chr13 | q34 | 111387781 | 1376 | | regulatory_region_variant | CpG: 15 | 14298 |
| Donor1_099 | chr13 | q33.3 | 108648141 | 2380 | MYO16 | intron_variant | CpG: 137 | 151470 |
| Donor1_100 | chr13 | q31.1 | 79765860 | 1406 | | intergenic_variant | CpG: 75 | 284282 |
| Donor1_101 | chr13 | q22.2 | 75498367 | 1314 | | intergenic_variant | CpG: 197 | 15704 |
| Donor1_102 | chr13 | q14.2 | 50045597 | 2442 | DLEU2 | intron_variant&non_coding_transcript_variant | CpG: 101 | 35549 |
| Donor1_103 | chr13 | q14.2 | 48970186 | 1114 | | downstream_gene_variant | CpG: 132 | 5501 |
| Donor1_104 | chr13 | q14.11 | 41769431 | 988 | VWA8 | intron_variant | CpG: 71 | 191314 |
| Donor1_105 | chr13 | q14.11 | 40983099 | 473 | ELF1 | intron_variant | CpG: 20 | 60429 |
| Donor1_106 | chr13 | q13.1 | 32239706 | 1070 | FRY | intron_variant | CpG: 23 | 71582 |
| Donor1_107 | chr13 | q12.12 | 24202990 | 2336 | | intron_variant&NMD_transcript_variant | CpG: 66 | 41591 |
| Donor1_108 | chr13 | q12.11 | 21345780 | 182 | LINC00539 | intron_variant&non_coding_transcript_variant | CpG: 27 | 19282 |
| Donor1_109 | chr14 | q32.2 | 96386860 | 1165 | GSKIP | 3_prime_UTR_variant | CpG: 43 | 4415 |
| Donor1_110 | chr14 | q32.12 | 92543045 | 1065 | RIN3 | intron_variant | CpG: 141 | 28161 |
| Donor1_111 | chr14 | q24.2 | 72468735 | 555 | RGS6 | intron_variant | CpG: 165 | 423347 |
| Donor1_112 | chr14 | q21.2 | 43203249 | 409 | | intergenic_variant | CpG: 47 | 1596123 |
| Donor1_113 | chr14 | q21.1 | 42144145 | 342 | | intergenic_variant | CpG: 47 | 537019 |
| Donor1_114 | chr14 | q21.1 | 41742281 | 618 | LRFN5 | intron_variant | CpG: 47 | 135155 |
| Donor1_115 | chr14 | q11.2 | 20622737 | 389 | | intergenic_variant | CpG: 27 | 199 |
| Donor1_116 | chr15 | q26.3 | 100624625 | 1870 | ASB7 | intron_variant | CpG: 80 | 21693 |
| Donor1_117 | chr15 | q25.3 | 84761328 | 1996 | ZNF592 | intron_variant | CpG: 151 | 12196 |
| Donor1_118 | chr15 | q25.3 | 85736401 | 927 | AKAP13 | intron_variant | CpG: 20 | 22318 |
| Donor1_119 | chr15 | q25.1 | 79433488 | 1336 | MINAR1 | intron_variant | CpG: 165 | 187 |
| Donor1_120 | chr15 | q24.3 | 77934935 | 943 | COMMD4P1 | downstream_gene_variant | CpG: 35 | 9412 |
| Donor1_121 | chr15 | q24.3 | 77934953 | 943 | COMMD4P1 | downstream_gene_variant | CpG: 35 | 9394 |
| Donor1_122 | chr15 | q22.2 | 61713988 | 1355 | | intron_variant&non_coding_transcript_variant | CpG: 83 | 346041 |
| Donor1_123 | chr15 | q21.1 | 46743199 | 1895 | | intron_variant&non_coding_transcript_variant | CpG: 96 | 440974 |
| Donor1_124 | chr15 | q21.1 | 47526479 | 951 | | intron_variant&non_coding_transcript_variant | CpG: 129 | 190743 |
| Donor1_125 | chr15 | q21.1 | 49066458 | 397 | | regulatory_region_variant | CpG: 43 | 19884 |
| Donor1_126 | chr15 | q15.3 | 44125831 | 1594 | FRMD5 | intron_variant | CpG: 105 | 68713 |
| Donor1_127 | chr15 | q14 | 36572624 | 267 | COX6CP4 | downstream_gene_variant | CpG: 35 | 6635 |
| Donor1_128 | chr16 | q22.1 | 67961195 | 731 | SLC12A4 | intron_variant | CpG: 79 | 6982 |
| Donor1_129 | chr16 | q22.1 | 67023248 | 16 | RN7SL543P | downstream_gene_variant | CpG: 143 | 5463 |
| Donor1_130 | chr16 | p13.3 | 2010173 | 2230 | ZNF598 | upstream_gene_variant | CpG: 102 | 142 |
| Donor1_131 | chr16 | p13.13 | 11728499 | 1133 | TXNDC11 | intron_variant | CpG: 114 | 13719 |
| Donor1_132 | chr16 | p12.3 | 17365628 | 2217 | XYLT1 | intron_variant | CpG: 103 | 104660 |
| Donor1_133 | chr16 | p12.2 | 23338952 | 1418 | SCNN1B | intron_variant | CpG: 62 | 36327 |
| Donor1_134 | chr16 | p12.2 | 21940496 | 958 | | intron_variant&non_coding_transcript_variant | CpG: 115 | 67171 |
| Donor1_135 | chr17 | q25.3 | 79792801 | 1752 | CBX8 | 3_prime_UTR_variant | CpG: 297 | 0 |
| Donor1_136 | chr17 | q25.3 | 81397241 | 858 | BAHCC1 | intron_variant | CpG: 653 | 0 |

| | | | | | | | | |
|---|---|---|---|---|---|---|---|---|
| Donor1_137 | chr17 | q25.3 | 81401011 | 550 | BAHCC1 | intron_variant | CpG: 653 | 69 |
| Donor1_138 | chr17 | q23.3 | 63583116 | 3294 | DCAF7 | intron_variant | CpG: 52 | 17680 |
| Donor1_139 | chr17 | q23.2 | 63006411 | 1883 | TANC2 | upstream_gene_variant | CpG: 145 | 39364 |
| Donor1_140 | chr17 | q22 | 55940396 | 1883 | ANKFN1 | intron_variant | CpG: 34 | 94618 |
| Donor1_141 | chr17 | q21.32 | 47127721 | 1765 | CDC27 | intron_variant | CpG: 45 | 27332 |
| Donor1_142 | chr17 | q21.31 | 45151470 | 909 | HEXIM1 | 3_prime_UTR_variant | CpG: 192 | 1221 |
| Donor1_143 | chr17 | q12 | 35516422 | 1233 | SLFN12L | intron_variant | CpG: 51 | 28494 |
| Donor1_144 | chr17 | q11.2 | 31596686 | 1233 | | intron_variant&non_coding_transcript_variant | CpG: 196 | 36328 |
| Donor1_145 | chr17 | p12 | 16086944 | 2860 | NCOR1 | intron_variant | CpG: 130 | 86570 |
| Donor1_146 | chr18 | q21.33 | 63294043 | 1220 | BCL2 | intron_variant | CpG: 236 | 24228 |
| Donor1_147 | chr18 | q21.31 | 57355405 | 1354 | ST8SIA3 | intron_variant | CpG: 154 | 1031 |
| Donor1_148 | chr18 | q21.2 | 54276498 | 572 | POLI | intron_variant | CpG: 52 | 6536 |
| Donor1_149 | chr18 | q12.2 | 35970303 | 478 | C18orf21 | upstream_gene_variant | CpG: 50 | 1886 |
| Donor1_150 | chr18 | p11.31 | 3103248 | 2037 | MYOM1 | intron_variant | CpG: 20 | 35713 |
| Donor1_151 | chr18 | p11.31 | 6215115 | 274 | L3MBTL4 | intron_variant | CpG: 22 | 69261 |
| Donor1_152 | chr19 | q13.43 | 56023170 | 1374 | NLRP5 | intron_variant | CpG: 178 | 63503 |
| Donor1_153 | chr19 | q13.43 | 56842021 | 239 | PEG3 | upstream_gene_variant | CpG: 83 | 1105 |
| Donor1_154 | chr19 | q13.42 | 53158478 | 24 | ZNF665 | downstream_gene_variant | CpG: 37 | 0 |
| Donor1_155 | chr19 | q13.41 | 52886449 | 3651 | ZNF320 | intron_variant | CpG: 56 | 10759 |
| Donor1_156 | chr19 | q13.41 | 52032129 | 2509 | ZNF432 | 3_prime_UTR_variant | CpG: 42 | 3658 |
| Donor1_157 | chr19 | q13.41 | 52815124 | 730 | ZNF28 | intron_variant | CpG: 46 | 28165 |
| Donor1_158 | chr19 | q13.32 | 46784888 | 2285 | SLC1A5 | intron_variant | CpG: 105 | 2441 |
| Donor1_159 | chr19 | q13.32 | 46784894 | 2285 | SLC1A5 | intron_variant | CpG: 105 | 2435 |
| Donor1_160 | chr19 | q13.2 | 41931485 | 1326 | ERFL | upstream_gene_variant | CpG: 38 | 3155 |
| Donor1_161 | chr19 | q13.12 | 37639601 | 2025 | ZFP30 | intron_variant | CpG: 119 | 15325 |
| Donor1_162 | chr19 | q13.12 | 36688272 | 1527 | ZNF567 | intron_variant | CpG: 53 | 585 |
| Donor1_163 | chr19 | q13.12 | 37487781 | 995 | ZNF570 | 3_prime_UTR_variant | CpG: 64 | 18068 |
| Donor1_164 | chr19 | q13.12 | 36685723 | 638 | ZNF567 | upstream_gene_variant | CpG: 53 | 1480 |
| Donor1_165 | chr19 | q13.12 | 37628578 | 408 | ZFP30 | downstream_gene_variant | CpG: 119 | 26348 |
| Donor1_166 | chr19 | p13.3 | 1903447 | 120 | ADAT3 | upstream_gene_variant | CpG: 62 | 1661 |
| Donor1_167 | chr19 | p13.3 | 3182569 | 18 | NCLN | upstream_gene_variant | CpG: 95 | 2581 |
| Donor1_168 | chr19 | p13.2 | 12152296 | 1722 | ZNF625 | intron_variant | CpG: 58 | 3888 |
| Donor1_169 | chr19 | p13.2 | 9640030 | 505 | ZNF562 | downstream_gene_variant | CpG: 22 | 18712 |
| Donor1_170 | chr19 | p13.2 | 12044087 | 48 | ZNF878 | coding_sequence_variant | CpG: 55 | 8253 |
| Donor1_171 | chr19 | p13.11 | 18515736 | 344 | ELL | intron_variant | CpG: 116 | 5753 |
| Donor1_172 | chr19 | p12 | 20556504 | 2890 | ZNF737 | intron_variant | CpG: 26 | 504206 |
| Donor1_173 | chr2 | q37.3 | 241697138 | 17 | ING5 | upstream_gene_variant | CpG: 167 | 4291 |
| Donor1_174 | chr2 | q37.1 | 230714280 | 969 | CAB39 | intron_variant | CpG: 125 | 491 |
| Donor1_175 | chr2 | q37.1 | 232315264 | 930 | DIS3L2 | intron_variant | CpG: 174 | 35707 |
| Donor1_176 | chr2 | q35 | 217015640 | 440 | LOC101928278 | intron_variant&non_coding_transcript_variant | CpG: 18 | 205149 |
| Donor1_177 | chr2 | q33.3 | 206145138 | 25 | NDUFS1 | intron_variant | CpG: 94 | 14049 |
| Donor1_178 | chr2 | q33.1 | 201073794 | 3318 | NDUFB3 | intron_variant | CpG: 29 | 2141 |
| Donor1_179 | chr2 | q32.1 | 183810384 | 486 | | intergenic_variant | CpG: 26 | 549529 |
| Donor1_180 | chr2 | q31.3 | 181459310 | 1429 | ITGA4 | intron_variant | CpG: 119 | 1008 |
| Donor1_181 | chr2 | q24.3 | 167876876 | 2108 | B3GALT1 | downstream_gene_variant | CpG: 26 | 160552 |
| Donor1_182 | chr2 | q24.3 | 168233051 | 1216 | STK39 | intron_variant | CpG: 128 | 13390 |
| Donor1_183 | chr2 | q21.2 | 133187695 | 926 | NCKAP5 | intron_variant | CpG: 52 | 78681 |
| Donor1_184 | chr2 | q13 | 110815548 | 128 | ACOXL | intron_variant | CpG: 89 | 82241 |
| Donor1_185 | chr2 | q11.2 | 98725866 | 3187 | MGAT4A | intron_variant | CpG: 150 | 4554 |
| Donor1_186 | chr2 | q11.2 | 98662812 | 527 | MGAT4A | intron_variant | CpG: 84 | 53953 |
| Donor1_187 | chr2 | p25.3 | 1156078 | 258 | SNTG2 | intron_variant | CpG: 50 | 11253 |
| Donor1_188 | chr2 | p25.1 | 9830430 | 1404 | | regulatory_region_variant | CpG: 76 | 12705 |
| Donor1_189 | chr2 | p24.3 | 15228607 | 1156 | NBAS | intron_variant | CpG: 27 | 332409 |
| Donor1_190 | chr2 | p24.2 | 16939068 | 551 | | intergenic_variant | CpG: 50 | 430514 |
| Donor1_191 | chr2 | p24.1 | 20673450 | 809 | GDF7 | 3_prime_UTR_variant | CpG: 143 | 1930 |
| Donor1_192 | chr2 | p23.3 | 24044817 | 1330 | FKBP1B | upstream_gene_variant | CpG: 59 | 5014 |
| Donor1_193 | chr2 | p22.3 | 33243906 | 42 | LTBP1 | intron_variant | CpG: 164 | 295809 |
| Donor1_194 | chr2 | p11.2 | 84793411 | 719 | DNAH6 | intron_variant | CpG: 77 | 87197 |
| Donor1_195 | chr20 | q13.33 | 62909506 | 2269 | DIDO1 | intron_variant | CpG: 39 | 1497 |
| Donor1_196 | chr20 | q13.33 | 63896259 | 997 | DNAJC5 | intron_variant | CpG: 113 | 573 |
| Donor1_197 | chr20 | q13.2 | 51500377 | 1305 | NFATC2 | intron_variant | CpG: 43 | 7459 |
| Donor1_198 | chr20 | q13.2 | 53149361 | 557 | TSHZ2 | intron_variant | CpG: 27 | 175880 |
| Donor1_199 | chr20 | q13.12 | 44964057 | 699 | STK4 | upstream_gene_variant | CpG: 43 | 2197 |
| Donor1_200 | chr20 | q11.21 | 31559839 | 2269 | HM13 | intron_variant | CpG: 25 | 12350 |
| Donor1_201 | chr20 | q11.21 | 32424740 | 903 | ASXL1 | intron_variant | CpG: 22 | 31398 |
| Donor1_202 | chr20 | p13 | 833148 | 2718 | FAM110A | upstream_gene_variant | CpG: 63 | 330 |
| Donor1_203 | chr20 | p11.21 | 23541797 | 2523 | CST13P | non_coding_transcript_exon_variant | CpG: 87 | 95473 |
| Donor1_204 | chr21 | q22.11 | 31717650 | 756 | SCAF4 | intron_variant | CpG: 145 | 13494 |
| Donor1_205 | chr21 | q21.3 | 28183621 | 571 | LINC01695 | intron_variant&non_coding_transcript_variant | CpG: 29 | 701483 |
| Donor1_206 | chr21 | q21.2 | 25487689 | 1953 | | regulatory_region_variant | CpG: 51 | 74423 |

| | | | | | | | | |
|---|---|---|---|---|---|---|---|---|
| Donor1_207 | chr21 | q21.1 | 15241392 | 300 | | regulatory_region_variant | CpG: 171 | 175849 |
| Donor1_208 | chr21 | q21.1 | 17793595 | 925 | C21orf91 | intron_variant | CpG: 118 | 25184 |
| Donor1_209 | chr22 | q13.31 | 44378083 | 348 | | intergenic_variant | CpG: 154 | 46025 |
| Donor1_210 | chr22 | q13.1 | 39056087 | 465 | APOBEC3F | downstream_gene_variant | CpG: 107 | 88931 |
| Donor1_211 | chr3 | q29 | 197737339 | 2249 | RUBCN | upstream_gene_variant | CpG: 94 | 66 |
| Donor1_212 | chr3 | q29 | 197275776 | 34 | DLG1 | intron_variant | CpG: 115 | 21781 |
| Donor1_213 | chr3 | q26.32 | 177419415 | 745 | | intergenic_variant | CpG: 133 | 221349 |
| Donor1_214 | chr3 | q26.1 | 165773093 | 2434 | BCHE | 3_prime_UTR_variant | CpG: 23 | 1607047 |
| Donor1_215 | chr3 | q25.31 | 157133514 | 350 | LINC00881 | intron_variant&non_coding_transcript_variant | CpG: 46 | 12878 |
| Donor1_216 | chr3 | q25.1 | 152261499 | 1293 | MBNL1 | intron_variant&non_coding_transcript_variant | CpG: 92 | 7040 |
| Donor1_217 | chr3 | q22.3 | 136753312 | 862 | STAG1 | upstream_gene_variant | CpG: 172 | 231 |
| Donor1_218 | chr3 | q21.1 | 122689733 | 1347 | PARP14 | intron_variant | CpG: 50 | 8390 |
| Donor1_219 | chr3 | q21.1 | 123521846 | 663 | HACD2 | intron_variant | CpG: 87 | 62647 |
| Donor1_220 | chr3 | q13.2 | 112001867 | 1074 | TAGLN3 | intron_variant | CpG: 40 | 22571 |
| Donor1_221 | chr3 | q13.2 | 112542133 | 463 | ATG3 | intron_variant | CpG: 77 | 19043 |
| Donor1_222 | chr3 | q13.13 | 111571477 | 1175 | CD96 | intron_variant | CpG: 21 | 103004 |
| Donor1_223 | chr3 | p24.3 | 18679305 | 52 | SATB1-AS1 | intron_variant&non_coding_transcript_variant | CpG: 145 | 233764 |
| Donor1_224 | chr3 | p24.1 | 28462709 | 585 | ZCWPW2 | intron_variant | CpG: 111 | 112464 |
| Donor1_225 | chr3 | p21.33 | 43772398 | 312 | | intergenic_variant | CpG: 132 | 80755 |
| Donor1_226 | chr3 | p21.31 | 46041053 | 941 | XCR1 | intron_variant | CpG: 48 | 45072 |
| Donor1_227 | chr3 | p21.2 | 50789209 | 757 | DOCK3 | intron_variant | CpG: 133 | 113633 |
| Donor1_228 | chr3 | p13 | 71859228 | 1605 | | regulatory_region_variant | CpG: 74 | 73726 |
| Donor1_229 | chr3 | p13 | 71131047 | 976 | | intron_variant | CpG: 50 | 65377 |
| Donor1_230 | chr4 | q35.1 | 185560355 | 362 | | regulatory_region_variant | CpG: 85 | 24645 |
| Donor1_231 | chr4 | q34.2 | 176161953 | 3611 | WDR17 | intron_variant | CpG: 34 | 33440 |
| Donor1_232 | chr4 | q32.3 | 168482369 | 914 | DDX60L | upstream_gene_variant | CpG: 37 | 1795 |
| Donor1_233 | chr4 | q32.1 | 158810239 | 531 | FNIP2 | intron_variant | CpG: 85 | 40568 |
| Donor1_234 | chr4 | q31.3 | 154416070 | 1828 | DCHS2 | intron_variant | CpG: 24 | 675 |
| Donor1_235 | chr4 | q31.21 | 142417306 | 482 | INPP4B | intron_variant | CpG: 111 | 428482 |
| Donor1_236 | chr4 | q31.1 | 139283873 | 1422 | MGARP | upstream_gene_variant | CpG: 41 | 3578 |
| Donor1_237 | chr4 | q26 | 118279810 | 1706 | PRSS12 | downstream_gene_variant | CpG: 53 | 772 |
| Donor1_238 | chr4 | q25 | 108353757 | 2288 | | intergenic_variant | CpG: 136 | 180367 |
| Donor1_239 | chr4 | q23 | 99686605 | 1903 | | intergenic_variant | CpG: 20 | 33108 |
| Donor1_240 | chr4 | q23 | 99735186 | 1423 | | intergenic_variant | CpG: 20 | 81689 |
| Donor1_241 | chr4 | q22.3 | 94346062 | 1982 | HPGDS | upstream_gene_variant | CpG: 88 | 105589 |
| Donor1_242 | chr4 | q22.1 | 87962237 | 529 | | regulatory_region_variant | CpG: 135 | 45092 |
| Donor1_243 | chr4 | q22.1 | 88074591 | 118 | PKD2 | intron_variant | CpG: 135 | 66084 |
| Donor1_244 | chr4 | q21.1 | 77088847 | 1805 | | intergenic_variant | CpG: 106 | 12429 |
| Donor1_245 | chr4 | p16.3 | 372312 | 1236 | ZNF141 | intron_variant | CpG: 38 | 13799 |
| Donor1_246 | chr4 | p16.1 | 6701441 | 758 | S100P | downstream_gene_variant | CpG: 72 | 7781 |
| Donor1_247 | chr4 | p16.1 | 6701442 | 758 | S100P | downstream_gene_variant | CpG: 72 | 7780 |
| Donor1_248 | chr4 | p15.32 | 17534166 | 1307 | | intergenic_variant | CpG: 79 | 21808 |
| Donor1_249 | chr4 | p14 | 40204235 | 2818 | RHOH | intron_variant | CpG: 159 | 146971 |
| Donor1_250 | chr4 | p14 | 39574006 | 435 | SMIM14 | intron_variant | CpG: 99 | 45903 |
| Donor1_251 | chr5 | q33.2 | 156016683 | 1247 | SGCD | intron_variant | CpG: 125 | 287309 |
| Donor1_252 | chr5 | q33.2 | 154758047 | 699 | LARP1 | intron_variant | CpG: 302 | 598 |
| Donor1_253 | chr5 | q31.3 | 140450815 | 737 | ANKHD1 | intron_variant | CpG: 143 | 48292 |
| Donor1_254 | chr5 | q31.2 | 139850126 | 2874 | NRG2 | intron_variant | CpG: 98 | 1424 |
| Donor1_255 | chr5 | q31.1 | 131641683 | 26 | | intron_variant | CpG: 85 | 5871 |
| Donor1_256 | chr5 | q14.2 | 83074208 | 1002 | TMEM167A | intron_variant | CpG: 23 | 3076 |
| Donor1_257 | chr5 | q14.1 | 81857596 | 1206 | | regulatory_region_variant | CpG: 217 | 105566 |
| Donor1_258 | chr5 | q13.2 | 73450702 | 48 | FOXD1 | upstream_gene_variant | CpG: 66 | 463 |
| Donor1_259 | chr5 | q12.1 | 61020176 | 3218 | NDUFAF2 | intron_variant | CpG: 56 | 74513 |
| Donor1_260 | chr5 | q11.2 | 52987740 | 2189 | ITGA2 | upstream_gene_variant | CpG: 63 | 1146 |
| Donor1_261 | chr5 | q11.2 | 59130947 | 705 | PDE4D | intron_variant | CpG: 120 | 90893 |
| Donor1_262 | chr5 | q11.2 | 55174688 | 494 | CDC20B | upstream_gene_variant | CpG: 47 | 1192 |
| Donor1_263 | chr5 | p15.33 | 254611 | 38 | SDHA | intron_variant | CpG: 275 | 4076 |
| Donor1_264 | chr5 | p14.3 | 20682727 | 3717 | LINC02241 | intron_variant&non_coding_transcript_variant | CpG: 45 | 376900 |
| Donor1_265 | chr5 | p13.3 | 32536719 | 115 | SUB1 | intron_variant | CpG: 89 | 48779 |
| Donor1_266 | chr5 | p12 | 42876256 | 1567 | | intron_variant&non_coding_transcript_variant | CpG: 47 | 47857 |
| Donor1_267 | chr6 | q25.1 | 149351092 | 47 | TAB2 | intron_variant | CpG: 113 | 32710 |
| Donor1_268 | chr6 | q24.1 | 139325199 | 1302 | | intron_variant&non_coding_transcript_variant | CpG: 18 | 47884 |
| Donor1_269 | chr6 | q23.3 | 135177499 | 738 | MYB | upstream_gene_variant | CpG: 216 | 3352 |
| Donor1_270 | chr6 | q23.3 | 137918376 | 611 | | intergenic_variant | CpG: 82 | 50372 |
| Donor1_271 | chr6 | q22.31 | 119074936 | 2838 | FAM184A | intron_variant | CpG: 127 | 2991 |
| Donor1_272 | chr6 | q21 | 108874499 | 854 | ARMC2 | intron_variant | CpG: 62 | 25599 |
| Donor1_273 | chr6 | q21 | 106517358 | 726 | CRYBG1 | intron_variant | CpG: 119 | 4248 |
| Donor1_274 | chr6 | q21 | 109125168 | 56 | CEP57L1 | intron_variant | CpG: 111 | 29624 |
| Donor1_275 | chr6 | q21 | 106517559 | 1 | CRYBG1 | intron_variant | CpG: 119 | 4449 |
| Donor1_276 | chr6 | q16.2 | 99800755 | 98 | | intergenic_variant | CpG: 75 | 181354 |

| | | | | | | | | |
|---|---|---|---|---|---|---|---|---|
| Donor1_277 | chr6 | q15 | 87330259 | 1873 | SMIM8 | intron_variant | CpG: 30 | 7548 |
| Donor1_278 | chr6 | q14.1 | 80171134 | 3785 | BCKDHB | intron_variant | CpG: 53 | 64165 |
| Donor1_279 | chr6 | q14.1 | 75278465 | 877 | TMEM30A | intron_variant | CpG: 63 | 5823 |
| Donor1_280 | chr6 | q13 | 73765089 | 2948 | CD109 | intron_variant | CpG: 50 | 68678 |
| Donor1_281 | chr6 | p25.3 | 1770766 | 478 | GMDS | intron_variant | CpG: 105 | 145533 |
| Donor1_282 | chr6 | p25.1 | 6786667 | 214 | LOC101928004 | intron_variant&non_coding_transcript_variant | CpG: 72 | 239670 |
| Donor1_283 | chr6 | p24.3 | 10585184 | 654 | GCNT2 | intron_variant | CpG: 56 | 109201 |
| Donor1_284 | chr6 | p24.3 | 8181400 | 27 | | intron_variant&non_coding_transcript_variant | CpG: 38 | 78701 |
| Donor1_285 | chr6 | p22.3 | 15837722 | 4217 | LOC105374948 | upstream_gene_variant | CpG: 95 | 174541 |
| Donor1_286 | chr6 | p22.3 | 24667496 | 1312 | ACOT13 | intron_variant | CpG: 76 | 329 |
| Donor1_287 | chr6 | p22.3 | 24893120 | 1213 | RIPOR2 | intron_variant | CpG: 82 | 17279 |
| Donor1_288 | chr6 | p22.3 | 24959334 | 1112 | RIPOR2 | intron_variant | CpG: 82 | 47976 |
| Donor1_289 | chr6 | p22.3 | 25133259 | 1082 | CMAHP | intron_variant&non_coding_transcript_variant | CpG: 36 | 6434 |
| Donor1_290 | chr6 | p22.1 | 27167762 | 3159 | | intergenic_variant | CpG: 29 | 9826 |
| Donor1_291 | chr6 | p21.1 | 43241923 | 1960 | TTBK1 | upstream_gene_variant | CpG: 27 | 1454 |
| Donor1_292 | chr6 | p21.1 | 42448770 | 1559 | TRERF1 | intron_variant | CpG: 18 | 3003 |
| Donor1_293 | chr6 | p21.1 | 43142488 | 693 | PTK7 | intron_variant | CpG: 159 | 28486 |
| Donor1_294 | chr7 | q36.1 | 150553002 | 970 | LOC107986860 | intron_variant | CpG: 96 | 144560 |
| Donor1_295 | chr7 | q36.1 | 148793223 | 418 | CUL1 | intron_variant | CpG: 34 | 9811 |
| Donor1_296 | chr7 | q34 | 138756921 | 1198 | ATP6V0A4 | intron_variant | CpG: 74 | 91801 |
| Donor1_297 | chr7 | q33 | 134737920 | 3292 | TUBB3P2 | upstream_gene_variant | CpG: 63 | 248449 |
| Donor1_298 | chr7 | q33 | 134315507 | 2055 | SLC35B4 | intron_variant | CpG: 75 | 949 |
| Donor1_299 | chr7 | q33 | 134449197 | 871 | AKR1B1 | intron_variant | CpG: 93 | 9167 |
| Donor1_300 | chr7 | q31.31 | 121354118 | 996 | FAM3C | intron_variant | CpG: 81 | 23429 |
| Donor1_301 | chr7 | q31.2 | 115875652 | 706 | | intergenic_variant | CpG: 71 | 334665 |
| Donor1_302 | chr7 | q31.1 | 110987573 | 1273 | IMMP2L | intron_variant | CpG: 68 | 574451 |
| Donor1_303 | chr7 | q31.1 | 107924819 | 638 | LAMB1 | intron_variant | CpG: 51 | 33281 |
| Donor1_304 | chr7 | q21.2 | 91606109 | 1175 | | intergenic_variant | CpG: 35 | 274575 |
| Donor1_305 | chr7 | q21.11 | 80116445 | 1606 | GNAI1 | intron_variant | CpG: 135 | 18033 |
| Donor1_306 | chr7 | q11.23 | 76356558 | 2741 | YWHAG | intron_variant | CpG: 97 | 1732 |
| Donor1_307 | chr7 | q11.23 | 74074120 | 1013 | ELN | downstream_gene_variant | CpG: 96 | 9395 |
| Donor1_308 | chr7 | p21.3 | 12696127 | 1678 | | downstream_gene_variant | CpG: 111 | 8504 |
| Donor1_309 | chr7 | p21.2 | 16113361 | 511 | CRPPA | intron_variant | CpG: 17 | 286137 |
| Donor1_310 | chr7 | p21.1 | 19512343 | 3058 | | intron_variant&non_coding_transcript_variant | CpG: 44 | 196432 |
| Donor1_311 | chr7 | p21.1 | 19945274 | 1409 | | intron_variant&non_coding_transcript_variant | CpG: 44 | 236036 |
| Donor1_312 | chr7 | p14.3 | 32502739 | 1742 | AVL9 | intron_variant | CpG: 49 | 6811 |
| Donor1_313 | chr7 | p14.1 | 40046053 | 2245 | CDK13 | intron_variant | CpG: 109 | 88230 |
| Donor1_314 | chr8 | q24.21 | 126812846 | 892 | | intron_variant&non_coding_transcript_variant | CpG: 180 | 254321 |
| Donor1_315 | chr8 | q24.21 | 130124487 | 71 | ASAP1 | intron_variant | CpG: 136 | 107638 |
| Donor1_316 | chr8 | q24.13 | 123891294 | 23 | FER1L6 | intron_variant | CpG: 91 | 122269 |
| Donor1_317 | chr8 | q22.1 | 97645888 | 1255 | MTDH | intron_variant | CpG: 127 | 886 |
| Donor1_318 | chr8 | q12.2 | 60952337 | 1031 | | intron_variant&non_coding_transcript_variant | CpG: 83 | 29025 |
| Donor1_319 | chr8 | p12 | 30618570 | 1778 | GTF2E2 | intron_variant | CpG: 72 | 39084 |
| Donor1_320 | chr8 | p12 | 29535057 | 91 | | downstream_gene_variant | CpG: 28 | 181773 |
| Donor1_321 | chr9 | q33.3 | 126369456 | 1213 | MVB12B | intron_variant | CpG: 17 | 31161 |
| Donor1_322 | chr9 | q33.3 | 126369457 | 1213 | MVB12B | intron_variant | CpG: 17 | 31162 |
| Donor1_323 | chr9 | q32 | 112316004 | 799 | PTBP3 | intron_variant | CpG: 141 | 16866 |
| Donor1_324 | chr9 | q22.33 | 96992289 | 2492 | MFSD14C | intron_variant&non_coding_transcript_variant | CpG: 104 | 20528 |
| Donor1_325 | chr9 | q22.31 | 91207580 | 1596 | | non_coding_transcript_exon_variant | CpG: 76 | 13442 |
| Donor1_326 | chr9 | q22.2 | 89403769 | 819 | SEMA4D | intron_variant | CpG: 21 | 22454 |
| Donor1_327 | chr9 | q21.13 | 71938159 | 1390 | C9orf85 | intron_variant | CpG: 21 | 26396 |
| Donor1_328 | chr9 | p24.3 | 295159 | 3248 | DOCK8 | intron_variant | CpG: 101 | 79728 |
| Donor1_329 | chr9 | p24.3 | 295160 | 3248 | DOCK8 | intron_variant | CpG: 101 | 79729 |
| Donor1_330 | chr9 | p24.2 | 4303993 | 1531 | GLIS3 | upstream_gene_variant | CpG: 205 | 3811 |
| Donor1_331 | chr9 | p21.3 | 20387525 | 762 | MLLT3 | intron_variant | CpG: 167 | 232901 |
| Donor1_332 | chr9 | p21.1 | 28670688 | 958 | LINGO2 | upstream_gene_variant | CpG: 74 | 541486 |
| Donor1_333 | chr9 | p21.1 | 31976951 | 1439 | | intergenic_variant | CpG: 59 | 407472 |
| Donor1_334 | chrX | q28 | 153614613 | 2818 | | downstream_gene_variant | CpG: 32 | 3923 |
| Donor1_335 | chrX | q27.1 | 140108765 | 16 | | intergenic_variant | CpG: 55 | 17220 |
| Donor1_336 | chrX | q24 | 118554617 | 343 | DOCK11 | intron_variant | CpG: 77 | 58111 |
| Donor1_337 | chrX | q23 | 114022817 | 276 | XACT | intron_variant&non_coding_transcript_variant | CpG: 18 | 367681 |
| Donor1_338 | chrX | q21.1 | 77760742 | 755 | ATRX | intron_variant | CpG: 54 | 24776 |
| Donor1_339 | chrX | q13.1 | 68506983 | 1229 | YIPF6 | intron_variant | CpG: 57 | 7754 |
| Donor1_340 | chrX | p22.31 | 7926340 | 2320 | PNPLA4 | intron_variant | CpG: 49 | 697 |
| Donor1_341 | chrX | p22.13 | 17670318 | 938 | NHS | intron_variant | CpG: 84 | 13955 |
| Donor1_342 | chrX | p21.1 | 33595270 | 2374 | | intergenic_variant | CpG: 36 | 131044 |
| Donor1_343 | chrX | p11.3 | 47129123 | 656 | LOC107985720 | non_coding_transcript_exon_variant | CpG: 102 | 15229 |
| Donor1_344 | chrX | p11.3 | 42991657 | 1748 | | regulatory_region_variant | CpG: 34 | 213401 |
| Donor2_001 | chr1 | q42.3 | 236232986 | 301 | ERO1B | intron_variant | CpG: 108 | 48318 |
| Donor2_002 | chr1 | q42.12 | 226547896 | 570 | STUM | upstream_gene_variant | CpG: 120 | 784 |

| | | | | | | | | |
|---|---|---|---|---|---|---|---|---|
| Donor2_003 | chr1 | q41 | 220754962 | 304 | MTARC2 | coding_sequence_variant | CpG: 45 | 6407 |
| Donor2_004 | chr1 | q41 | 216703428 | 584 | ESRRG | intron_variant | CpG: 43 | 102036 |
| Donor2_005 | chr1 | q32.1 | 205616200 | 1296 | ELK4 | 3_prime_UTR_variant | CpG: 115 | 15166 |
| Donor2_006 | chr1 | q31.3 | 197863535 | 1072 | | intergenic_variant | CpG: 83 | 38924 |
| Donor2_007 | chr1 | p31.1 | 82848429 | 227 | | upstream_gene_variant | CpG: 50 | 1012156 |
| Donor2_008 | chr1 | p31.3 | 62688811 | 388 | | intron_variant&non_coding_transcript_variant | CpG: 111 | 133 |
| Donor2_009 | chr1 | p34.2 | 43295705 | 984 | | intergenic_variant | CpG: 43 | 9147 |
| Donor2_010 | chr1 | p35.1 | 32518786 | 636 | | intron_variant&NMD_transcript_variant | CpG: 64 | 17824 |
| Donor2_011 | chr1 | q24.2 | 169950498 | 118 | KIFAP3 | intron_variant | CpG: 66 | 56682 |
| Donor2_012 | chr1 | q21.3 | 154188068 | 1401 | TPM3 | intron_variant | CpG: 49 | 4618 |
| Donor2_013 | chr1 | p21.3 | 96826338 | 710 | PTBP2 | downstream_gene_variant | CpG: 64 | 104075 |
| Donor2_014 | chr1 | p22.1 | 93286524 | 31 | CCDC18-AS1 | intron_variant&non_coding_transcript_variant | CpG: 66 | 59138 |
| Donor2_015 | chr1 | p22.2 | 89818952 | 998 | LRRC8D | upstream_gene_variant | CpG: 147 | 1679 |
| Donor2_016 | chr1 | p31.1 | 77166285 | 306 | PIGK | intron_variant | CpG: 23 | 53027 |
| Donor2_017 | chr1 | p32.1 | 58698784 | 17 | MYSM1 | intron_variant | CpG: 271 | 83006 |
| Donor2_018 | chr1 | p34.3 | 37928806 | 1267 | INPP5B | intron_variant | CpG: 53 | 2759 |
| Donor2_019 | chr1 | p35.3 | 28757561 | 339 | YTHDF2 | intron_variant | CpG: 26 | 17718 |
| Donor2_020 | chr1 | p35.3 | 28327409 | 268 | MED18 | upstream_gene_variant | CpG: 27 | 1446 |
| Donor2_021 | chr1 | p36.11 | 23868525 | 68 | FUCA1 | upstream_gene_variant | CpG: 84 | 25 |
| Donor2_022 | chr1 | p36.21 | 12515130 | 470 | VPS13D | downstream_gene_variant | CpG: 18 | 25192 |
| Donor2_023 | chr1 | p36.22 | 9693582 | 534 | PIK3CD | intron_variant | CpG: 102 | 3412 |
| Donor2_024 | chr1 | p36.22 | 9671505 | 941 | PIK3CD | intron_variant | CpG: 102 | 17733 |
| Donor2_025 | chr1 | p36.23 | 8874272 | 865 | ENO1 | intron_variant | CpG: 112 | 3983 |
| Donor2_026 | chr10 | q26.11 | 118797971 | 320 | | intron_variant&non_coding_transcript_variant | CpG: 108 | 42375 |
| Donor2_027 | chr10 | p13 | 16139950 | 14 | | intergenic_variant | CpG: 29 | 279547 |
| Donor2_028 | chr10 | q22.2 | 73464488 | 792 | PPP3CB | intron_variant | CpG: 45 | 30788 |
| Donor2_029 | chr10 | q21.3 | 68307912 | 2010 | PBLD | intron_variant | CpG: 128 | 23387 |
| Donor2_030 | chr10 | q11.23 | 50421526 | 409 | SGMS1 | intron_variant | CpG: 53 | 3104 |
| Donor2_031 | chr10 | p11.21 | 34370262 | 381 | PARD3 | intron_variant | CpG: 188 | 443935 |
| Donor2_032 | chr10 | p11.22 | 32515971 | 1121 | CCDC7 | intron_variant | CpG: 164 | 168244 |
| Donor2_033 | chr10 | p11.22 | 32347873 | 471 | EPC1 | upstream_gene_variant | CpG: 164 | 146 |
| Donor2_034 | chr10 | p12.2 | 22317154 | 740 | COMMD3 | intron_variant | CpG: 102 | 451 |
| Donor2_035 | chr10 | p12.33 | 17443122 | 1130 | ST8SIA6 | intron_variant | CpG: 103 | 10287 |
| Donor2_036 | chr10 | p13 | 17228133 | 94 | VIM | upstream_gene_variant | CpG: 195 | 299 |
| Donor2_037 | chr10 | p14 | 11631596 | 503 | LOC105376413 | downstream_gene_variant | CpG: 127 | 19633 |
| Donor2_038 | chr11 | q24.3 | 128653336 | 378 | | intron_variant&non_coding_transcript_variant | CpG: 30 | 33634 |
| Donor2_039 | chr11 | q22.2 | 102468502 | 137 | | intron_variant&non_coding_transcript_variant | CpG: 106 | 15471 |
| Donor2_040 | chr11 | q22.1 | 98293846 | 451 | | intergenic_variant | CpG: 34 | 726968 |
| Donor2_041 | chr11 | p11.12 | 49327838 | 127 | | intron_variant&non_coding_transcript_variant | CpG: 38 | 119350 |
| Donor2_042 | chr11 | q23.3 | 118251333 | 1119 | MPZL3 | intron_variant | CpG: 77 | 97854 |
| Donor2_043 | chr11 | q23.3 | 118243883 | 476 | MPZL3 | intron_variant | CpG: 77 | 90404 |
| Donor2_044 | chr11 | q21 | 95088579 | 572 | ENDOD1 | upstream_gene_variant | CpG: 101 | 855 |
| Donor2_045 | chr11 | q14.1 | 83217780 | 75 | ANKRD42 | intron_variant | CpG: 90 | 23162 |
| Donor2_046 | chr11 | q13.4 | 75324161 | 179 | ARRB1 | intron_variant | CpG: 97 | 27020 |
| Donor2_047 | chr11 | q13.4 | 74757019 | 1618 | RNF169 | intron_variant | CpG: 62 | 7506 |
| Donor2_048 | chr11 | q13.4 | 74748590 | 501 | RNF169 | upstream_gene_variant | CpG: 62 | 487 |
| Donor2_049 | chr11 | q12.3 | 62726033 | 1165 | HNRNPUL2 | intron_variant | CpG: 142 | 490 |
| Donor2_050 | chr11 | p13 | 36290357 | 570 | COMMD9 | upstream_gene_variant | CpG: 122 | 86047 |
| Donor2_051 | chr11 | p14.1 | 30685446 | 179 | LINC02859 | non_coding_transcript_exon_variant | CpG: 132 | 98865 |
| Donor2_052 | chr11 | p15.4 | 10859308 | 177 | ZBED5 | upstream_gene_variant | CpG: 97 | 402 |
| Donor2_053 | chr11 | p15.5 | 227938 | 1877 | SIRT3 | intron_variant | CpG: 101 | 7981 |
| Donor2_054 | chr12 | q22 | 94776619 | 1083 | | regulatory_region_variant | CpG: 119 | 96438 |
| Donor2_055 | chr12 | q24.33 | 133033326 | 768 | ZNF84 | upstream_gene_variant | CpG: 84 | 3854 |
| Donor2_056 | chr12 | q24.33 | 133010316 | 1167 | ZNF26 | coding_sequence_variant | CpG: 72 | 23449 |
| Donor2_057 | chr12 | q15 | 68586791 | 1607 | | intergenic_variant | CpG: 132 | 23650 |
| Donor2_058 | chr12 | q24.33 | 128823149 | 207 | SLC15A4 | intron_variant | CpG: 75 | 241 |
| Donor2_059 | chr12 | p13.31 | 6425349 | 1018 | | intergenic_variant | CpG: 55 | 40993 |
| Donor2_060 | chr12 | q24.31 | 123634635 | 309 | GTF2H3 | intron_variant | CpG: 43 | 703 |
| Donor2_061 | chr12 | q24.23 | 117920586 | 564 | KSR2 | intron_variant | CpG: 49 | 48472 |
| Donor2_062 | chr12 | q23.3 | 105172630 | 1316 | APPL2 | downstream_gene_variant | CpG: 69 | 63090 |
| Donor2_063 | chr12 | q15 | 70261792 | 213 | CNOT2 | intron_variant | CpG: 102 | 17823 |
| Donor2_064 | chr12 | q15 | 70253158 | 262 | CNOT2 | intron_variant | CpG: 102 | 9189 |
| Donor2_065 | chr12 | q14.1 | 62351763 | 785 | USP15 | intron_variant | CpG: 74 | 90867 |
| Donor2_066 | chr12 | q13.2 | 54504639 | 957 | NCKAP1L | intron_variant | CpG: 66 | 83511 |
| Donor2_067 | chr12 | q13.11 | 47194896 | 406 | PCED1B | intron_variant | CpG: 44 | 40356 |
| Donor2_068 | chr12 | p12.3 | 16024714 | 464 | DERA | intron_variant | CpG: 54 | 113114 |
| Donor2_069 | chr12 | p13.31 | 6444232 | 440 | CD27 | upstream_gene_variant | CpG: 76 | 26158 |
| Donor2_070 | chr12 | p13.33 | 390092 | 12 | KDM5A | upstream_gene_variant | CpG: 37 | 721 |
| Donor2_071 | chr13 | q34 | 114036269 | 483 | RASA3 | intron_variant | CpG: 24 | 1872 |
| Donor2_072 | chr13 | q33.3 | 108216028 | 303 | LIG4 | upstream_gene_variant | CpG: 19 | 1012 |

| | | | | | | | | |
|---|---|---|---|---|---|---|---|---|
| Donor2_073 | chr13 | q22.1 | 74053768 | 706 | KLF12 | intron_variant | CpG: 31 | 80205 |
| Donor2_074 | chr13 | q14.11 | 43449445 | 2407 | ENOX1 | intron_variant | CpG: 159 | 336454 |
| Donor2_075 | chr13 | q12.11 | 20710830 | 225 | IL17D | intron_variant | CpG: 156 | 6553 |
| Donor2_076 | chr14 | q23.1 | 59394212 | 1108 | | intergenic_variant | CpG: 189 | 69705 |
| Donor2_077 | chr14 | q12 | 24276890 | 608 | | intron_variant&non_coding_transcript_variant | CpG: 51 | 5154 |
| Donor2_078 | chr14 | q24.3 | 75283708 | 952 | FOS | downstream_gene_variant | CpG: 233 | 3613 |
| Donor2_079 | chr14 | q24.1 | 68786403 | 313 | ZFP36L1 | downstream_gene_variant | CpG: 34 | 3557 |
| Donor2_080 | chr14 | q21.2 | 44926974 | 821 | KLHL28 | 3_prime_UTR_variant | CpG: 66 | 29069 |
| Donor2_081 | chr14 | q21.1 | 39434059 | 1008 | FBXO33 | upstream_gene_variant | CpG: 127 | 1057 |
| Donor2_082 | chr14 | q12 | 31587112 | 389 | NUBPL | intron_variant | CpG: 61 | 129508 |
| Donor2_083 | chr15 | q22.2 | 60403065 | 9 | | upstream_gene_variant | CpG: 19 | 4334 |
| Donor2_084 | chr15 | q22.2 | 63324849 | 323 | CA12 | 3_prime_UTR_variant | CpG: 32 | 2963 |
| Donor2_085 | chr15 | q21.3 | 58704113 | 520 | ADAM10 | intron_variant | CpG: 125 | 44770 |
| Donor2_086 | chr15 | q21.3 | 55268761 | 4180 | RAB27A | intron_variant | CpG: 88 | 20814 |
| Donor2_087 | chr15 | q21.2 | 52045234 | 239 | MAPK6 | intron_variant | CpG: 117 | 25251 |
| Donor2_088 | chr15 | q15.1 | 41801915 | 902 | MAPKBP1 | intron_variant | CpG: 92 | 25490 |
| Donor2_089 | chr16 | q23.2 | 81305225 | 1913 | | intergenic_variant | CpG: 141 | 9521 |
| Donor2_090 | chr16 | q23.2 | 81319995 | 14 | GAN | intron_variant | CpG: 141 | 4074 |
| Donor2_091 | chr16 | q21 | 58259822 | 56 | CCDC113 | intron_variant | CpG: 41 | 9642 |
| Donor2_092 | chr16 | q12.2 | 53061243 | 661 | CHD9 | intron_variant | CpG: 54 | 5876 |
| Donor2_093 | chr16 | q12.2 | 53061242 | 661 | CHD9 | intron_variant | CpG: 54 | 5875 |
| Donor2_094 | chr16 | p12.2 | 23647831 | 467 | DCTN5 | intron_variant | CpG: 51 | 6234 |
| Donor2_095 | chr16 | p13.13 | 12082217 | 177 | SNX29 | intron_variant | CpG: 64 | 34582 |
| Donor2_096 | chr17 | q25.3 | 79604031 | 618 | | upstream_gene_variant | CpG: 231 | 4737 |
| Donor2_097 | chr17 | q24.1 | 65523553 | 752 | | regulatory_region_variant | CpG: 66 | 13285 |
| Donor2_098 | chr17 | q12 | 35459422 | 757 | | downstream_gene_variant | CpG: 27 | 961 |
| Donor2_099 | chr17 | q25.1 | 74760372 | 1268 | SLC9A3R1 | intron_variant | CpG: 152 | 10559 |
| Donor2_100 | chr17 | q25.1 | 74750448 | 522 | SLC9A3R1 | intron_variant | CpG: 152 | 635 |
| Donor2_101 | chr17 | q25.1 | 74533991 | 946 | CD300LB | upstream_gene_variant | CpG: 56 | 79513 |
| Donor2_102 | chr17 | q24.2 | 67848997 | 560 | BPTF | intron_variant | CpG: 104 | 23075 |
| Donor2_103 | chr17 | q24.2 | 66495374 | 696 | PRKCA | intron_variant | CpG: 183 | 191875 |
| Donor2_104 | chr17 | q24.2 | 66373483 | 271 | PRKCA | intron_variant | CpG: 183 | 69984 |
| Donor2_105 | chr17 | q21.31 | 43656009 | 803 | MEOX1 | intron_variant | CpG: 72 | 9551 |
| Donor2_106 | chr17 | p13.1 | 7189590 | 660 | DLG4 | 3_prime_UTR_variant | CpG: 68 | 15021 |
| Donor2_107 | chr17 | p13.1 | 6655655 | 1351 | MED31 | upstream_gene_variant | CpG: 104 | 339 |
| Donor2_108 | chr17 | p13.3 | 2451781 | 46 | METTL16 | intron_variant | CpG: 236 | 49591 |
| Donor2_109 | chr18 | q22.2 | 70419336 | 1227 | | regulatory_region_variant | CpG: 21 | 11235 |
| Donor2_110 | chr18 | q22.3 | 73243225 | 52 | LINC02864 | intron_variant&non_coding_transcript_variant | CpG: 275 | 373589 |
| Donor2_111 | chr18 | p11.31 | 5133888 | 768 | BOD1P2 | upstream_gene_variant | CpG: 80 | 62357 |
| Donor2_112 | chr19 | q13.2 | 40107147 | 170 | | intergenic_variant | CpG: 26 | 16191 |
| Donor2_113 | chr19 | q13.12 | 36504193 | 50 | | intergenic_variant | CpG: 44 | 14362 |
| Donor2_114 | chr19 | p12 | 23990836 | 1549 | | intergenic_variant | CpG: 32 | 18879 |
| Donor2_115 | chr19 | p13.2 | 9731045 | 136 | | non_coding_transcript_exon_variant | CpG: 22 | 4637 |
| Donor2_116 | chr19 | q13.43 | 58142402 | 406 | ZNF329 | intron_variant | CpG: 62 | 7969 |
| Donor2_117 | chr19 | q13.43 | 57792352 | 1276 | ZNF586 | intron_variant | CpG: 62 | 5602 |
| Donor2_118 | chr19 | q13.42 | 53552984 | 843 | ZNF331 | intron_variant | CpG: 83 | 1177 |
| Donor2_119 | chr19 | q13.32 | 46957641 | 503 | ARHGAP35 | intron_variant | CpG: 40 | 46409 |
| Donor2_120 | chr19 | q13.32 | 45834666 | 329 | SYMPK | intron_variant | CpG: 62 | 18663 |
| Donor2_121 | chr19 | q13.12 | 37467599 | 820 | ZNF569 | upstream_gene_variant | CpG: 57 | 111 |
| Donor2_122 | chr19 | q13.12 | 36812740 | 236 | ZNF790 | downstream_gene_variant | CpG: 40 | 14919 |
| Donor2_123 | chr19 | q13.12 | 36638494 | 606 | ZNF461 | 3_prime_UTR_variant | CpG: 39 | 28237 |
| Donor2_124 | chr19 | q13.12 | 35236059 | 983 | FAM187B2P | upstream_gene_variant | CpG: 34 | 10842 |
| Donor2_125 | chr19 | q13.11 | 34309052 | 1425 | GARRE1 | intron_variant | CpG: 56 | 50115 |
| Donor2_126 | chr19 | p13.11 | 18152054 | 48 | PIK3R2 | upstream_gene_variant | CpG: 119 | 324 |
| Donor2_127 | chr19 | p13.12 | 15505671 | 772 | CYP4F22 | upstream_gene_variant | CpG: 27 | 33577 |
| Donor2_128 | chr19 | p13.2 | 12288184 | 555 | ZNF44 | intron_variant | CpG: 26 | 6328 |
| Donor2_129 | chr19 | p13.2 | 11096329 | 457 | LDLR | intron_variant | CpG: 66 | 5319 |
| Donor2_130 | chr19 | p13.2 | 9185323 | 2202 | OR7D2 | intron_variant | CpG: 70 | 24081 |
| Donor2_131 | chr2 | q37.3 | 241098504 | 526 | MTERF4 | intron_variant | CpG: 59 | 3625 |
| Donor2_132 | chr2 | q37.1 | 233683381 | 888 | UGT1A9 | intron_variant | CpG: 42 | 60220 |
| Donor2_133 | chr2 | q36.2 | 224766753 | 187 | DOCK10 | intron_variant | CpG: 135 | 180982 |
| Donor2_134 | chr2 | q35 | 218899575 | 510 | LINC01494 | downstream_gene_variant | CpG: 44 | 760 |
| Donor2_135 | chr2 | q35 | 218799733 | 410 | CYP27A1 | intron_variant | CpG: 70 | 17275 |
| Donor2_136 | chr2 | q33.1 | 200881424 | 319 | PPIL3 | coding_sequence_variant | CpG: 85 | 16570 |
| Donor2_137 | chr2 | q32.3 | 196161899 | 874 | STK17B | intron_variant | CpG: 81 | 8967 |
| Donor2_138 | chr2 | q32.2 | 191084448 | 67 | STAT4 | intron_variant | CpG: 55 | 63757 |
| Donor2_139 | chr2 | q32.2 | 190980829 | 474 | STAT1 | intron_variant | CpG: 70 | 33054 |
| Donor2_140 | chr2 | q31.1 | 169706121 | 65 | PHOSPHO2 | downstream_gene_variant | CpG: 59 | 11260 |
| Donor2_141 | chr2 | q24.2 | 160303591 | 860 | RBMS1 | intron_variant | CpG: 48 | 33184 |
| Donor2_142 | chr2 | q24.2 | 159932143 | 394 | PLA2R1 | 3_prime_UTR_variant | CpG: 79 | 27202 |

| | | | | | | | | |
|---|---|---|---|---|---|---|---|---|
| Donor2_143 | chr2 | q24.1 | 157453426 | 796 | CYTIP | intron_variant | CpG: 31 | 143862 |
| Donor2_144 | chr2 | q21.2 | 134296445 | 745 | MGAT5 | intron_variant | CpG: 57 | 104289 |
| Donor2_145 | chr2 | q14.2 | 119996620 | 1326 | RPL27P7 | upstream_gene_variant | CpG: 102 | 16202 |
| Donor2_146 | chr2 | q12.2 | 105865717 | 246 | NCK2 | intron_variant | CpG: 78 | 15615 |
| Donor2_147 | chr2 | p11.2 | 85415866 | 663 | SH2D6 | upstream_gene_variant | CpG: 29 | 1730 |
| Donor2_148 | chr2 | q33.3 | 204530835 | 777 | | regulatory_region_variant | CpG: 83 | 14382 |
| Donor2_149 | chr2 | q33.2 | 203674255 | 1514 | | upstream_gene_variant | CpG: 65 | 138809 |
| Donor2_150 | chr2 | q23.3 | 150706921 | 1289 | | intergenic_variant | CpG: 118 | 220244 |
| Donor2_151 | chr2 | q14.3 | 126645214 | 77 | | intron_variant&non_coding_transcript_variant | CpG: 51 | 10907 |
| Donor2_152 | chr2 | p12 | 77896126 | 804 | | intron_variant&non_coding_transcript_variant | CpG: 32 | 1096941 |
| Donor2_153 | chr2 | p12 | 75360520 | 1734 | | intergenic_variant | CpG: 96 | 159616 |
| Donor2_154 | chr2 | p16.1 | 55533829 | 744 | CFAP36 | intron_variant | CpG: 23 | 13966 |
| Donor2_155 | chr2 | p16.2 | 53965084 | 364 | PSME4 | intron_variant | CpG: 81 | 5428 |
| Donor2_156 | chr2 | p22.1 | 39928013 | 820 | SLC8A1-AS1 | intron_variant&non_coding_transcript_variant | CpG: 53 | 148690 |
| Donor2_157 | chr2 | p22.3 | 32659374 | 14 | TTC27 | intron_variant | CpG: 20 | 31171 |
| Donor2_158 | chr2 | p23.3 | 23914924 | 228 | ATAD2B | intron_variant | CpG: 113 | 11463 |
| Donor2_159 | chr2 | p24.3 | 15304039 | 1009 | NBAS | intron_variant | CpG: 27 | 256977 |
| Donor2_160 | chr2 | p25.2 | 6834841 | 478 | NRIR | intron_variant&non_coding_transcript_variant | CpG: 137 | 30192 |
| Donor2_161 | chr20 | q13.2 | 53608884 | 422 | | intron_variant&non_coding_transcript_variant | CpG: 189 | 14063 |
| Donor2_162 | chr20 | q13.13 | 50818133 | 2528 | BCAS4 | intron_variant | CpG: 18 | 22468 |
| Donor2_163 | chr20 | q11.23 | 38998763 | 962 | DHX35 | intron_variant | CpG: 59 | 36222 |
| Donor2_164 | chr20 | q11.23 | 38998762 | 962 | DHX35 | intron_variant | CpG: 59 | 36221 |
| Donor2_165 | chr20 | q11.22 | 34918270 | 675 | ACSS2 | intron_variant | CpG: 63 | 37010 |
| Donor2_166 | chr20 | p12.1 | 12889963 | 868 | LINC01722 | intron_variant&non_coding_transcript_variant | CpG: 217 | 330061 |
| Donor2_167 | chr21 | q21.2 | 25338263 | 706 | | intron_variant&non_coding_transcript_variant | CpG: 28 | 23532 |
| Donor2_168 | chr21 | q22.2 | 39412439 | 2143 | LCA5L | intron_variant | CpG: 19 | 23536 |
| Donor2_169 | chr21 | q22.11 | 33566612 | 857 | SON | intron_variant | CpG: 86 | 21471 |
| Donor2_170 | chr21 | q22.11 | 32453875 | 478 | EVA1C | intron_variant | CpG: 23 | 23818 |
| Donor2_171 | chr21 | q21.3 | 25729396 | 669 | ATP5PF | intron_variant | CpG: 121 | 5108 |
| Donor2_172 | chr22 | q13.31 | 44493195 | 1635 | RTL6 | 3_prime_UTR_variant | CpG: 159 | 3319 |
| Donor2_173 | chr22 | q12.1 | 26734371 | 875 | MIATNB | intron_variant&non_coding_transcript_variant | CpG: 46 | 22364 |
| Donor2_174 | chr22 | q12.1 | 26654132 | 568 | MIAT | intron_variant&non_coding_transcript_variant | CpG: 51 | 2986 |
| Donor2_175 | chr22 | q11.21 | 19937034 | 1581 | TXNRD2 | intron_variant | CpG: 37 | 4540 |
| Donor2_176 | chr3 | q29 | 197961667 | 844 | IQCG | upstream_gene_variant | CpG: 51 | 1204 |
| Donor2_177 | chr3 | q29 | 197002330 | 2124 | MELTF | 3_prime_UTR_variant | CpG: 191 | 0 |
| Donor2_178 | chr3 | q26.31 | 172939176 | 1909 | SPATA16 | intron_variant | CpG: 41 | 188121 |
| Donor2_179 | chr3 | q25.1 | 151208969 | 161 | MED12L | intron_variant | CpG: 223 | 121588 |
| Donor2_180 | chr3 | q25.1 | 150968316 | 818 | CLRN1 | intron_variant | CpG: 223 | 116894 |
| Donor2_181 | chr3 | q23 | 142345114 | 273 | XRN1 | intron_variant | CpG: 71 | 102247 |
| Donor2_182 | chr3 | q23 | 141076789 | 143 | SPSB4 | intron_variant | CpG: 72 | 10150 |
| Donor2_183 | chr3 | q22.1 | 131738759 | 480 | CPNE4 | intron_variant | CpG: 23 | 211582 |
| Donor2_184 | chr3 | q13.32 | 119234880 | 1118 | B4GALT4 | intron_variant | CpG: 75 | 5613 |
| Donor2_185 | chr3 | q13.31 | 116133461 | 1125 | LSAMP | intron_variant | CpG: 72 | 348727 |
| Donor2_186 | chr3 | q13.12 | 108116029 | 332 | LOC105374031 | intron_variant&non_coding_transcript_variant | CpG: 130 | 24333 |
| Donor2_187 | chr3 | q13.12 | 106595470 | 1425 | LOC101929485 | intron_variant&non_coding_transcript_variant | CpG: 49 | 645093 |
| Donor2_188 | chr3 | q13.11 | 105724535 | 283 | CBLB | intron_variant | CpG: 84 | 143901 |
| Donor2_189 | chr3 | p12.3 | 79017110 | 1255 | ROBO1 | intron_variant | CpG: 49 | 1804 |
| Donor2_190 | chr3 | q29 | 197319476 | 836 | | intergenic_variant | CpG: 25 | 20242 |
| Donor2_191 | chr3 | q21.3 | 129374460 | 915 | | intergenic_variant | CpG: 26 | 30170 |
| Donor2_192 | chr3 | p22.1 | 39442560 | 395 | | intron_variant&non_coding_transcript_variant | CpG: 60 | 35407 |
| Donor2_193 | chr3 | p14.3 | 56862955 | 286 | ARHGEF3 | intron_variant | CpG: 57 | 60773 |
| Donor2_194 | chr3 | p21.31 | 47924328 | 49 | MAP4 | intron_variant | CpG: 33 | 74274 |
| Donor2_195 | chr3 | p22.2 | 37346172 | 91 | GOLGA4 | intron_variant | CpG: 79 | 102382 |
| Donor2_196 | chr3 | p22.3 | 32685002 | 359 | CNOT10 | upstream_gene_variant | CpG: 57 | 24 |
| Donor2_197 | chr3 | p24.3 | 18392499 | 754 | SATB1 | intron_variant | CpG: 19 | 33450 |
| Donor2_198 | chr3 | p25.2 | 12380416 | 1178 | PPARG | intron_variant | CpG: 154 | 91582 |
| Donor2_199 | chr4 | q34.3 | 181162549 | 1718 | LINC00290 | upstream_gene_variant | CpG: 20 | 978577 |
| Donor2_200 | chr4 | q31.22 | 146643303 | 1339 | POU4F2 | downstream_gene_variant | CpG: 208 | 2554 |
| Donor2_201 | chr4 | q28.3 | 138242378 | 1081 | SLC7A11 | upstream_gene_variant | CpG: 21 | 318203 |
| Donor2_202 | chr4 | q24 | 101289746 | 1821 | PPP3CA | intron_variant | CpG: 178 | 56733 |
| Donor2_203 | chr4 | q22.1 | 91064007 | 1470 | CCSER1 | intron_variant | CpG: 22 | 224891 |
| Donor2_204 | chr4 | q13.3 | 70741427 | 738 | RUFY3 | intron_variant | CpG: 167 | 35518 |
| Donor2_205 | chr4 | q34.2 | 175569287 | 464 | | intergenic_variant | CpG: 26 | 432036 |
| Donor2_206 | chr4 | q31.21 | 144344869 | 1075 | | intron_variant&NMD_transcript_variant | CpG: 91 | 300222 |
| Donor2_207 | chr4 | q25 | 112117418 | 700 | | upstream_gene_variant | CpG: 120 | 27897 |
| Donor2_208 | chr4 | q25 | 110852017 | 561 | | intergenic_variant | CpG: 18 | 212365 |
| Donor2_209 | chr4 | p15.32 | 17092960 | 93 | | intergenic_variant | CpG: 79 | 418727 |
| Donor2_210 | chr4 | p14 | 40874942 | 92 | APBB2 | intron_variant | CpG: 40 | 17652 |
| Donor2_211 | chr4 | p15.33 | 13206103 | 75 | LOC105374494 | intron_variant&non_coding_transcript_variant | CpG: 97 | 277796 |
| Donor2_212 | chr4 | p16.1 | 9972016 | 1670 | SLC2A9 | intron_variant | CpG: 49 | 46842 |

| | | | | | | | | |
|---|---|---|---|---|---|---|---|---|
| Donor2_213 | chr5 | q35.3 | 180016649 | 318 | RNF130 | intron_variant | CpG: 134 | 54751 |
| Donor2_214 | chr5 | q35.3 | 179551340 | 892 | RUFY1 | intron_variant | CpG: 100 | 50 |
| Donor2_215 | chr5 | q33.3 | 158853398 | 72 | EBF1 | intron_variant | CpG: 25 | 197973 |
| Donor2_216 | chr5 | q31.3 | 141360650 | 880 | PCDHGB2 | coding_sequence_variant | CpG: 65 | 958 |
| Donor2_217 | chr5 | q23.2 | 126797368 | 925 | LMNB1 | intron_variant | CpG: 202 | 18831 |
| Donor2_218 | chr5 | q23.1 | 119318231 | 1061 | TNFAIP8 | intron_variant | CpG: 42 | 37476 |
| Donor2_219 | chr5 | q23.1 | 119275075 | 1075 | TNFAIP8 | intron_variant | CpG: 43 | 5936 |
| Donor2_220 | chr5 | q22.1 | 111435114 | 1437 | CAMK4 | intron_variant | CpG: 51 | 77009 |
| Donor2_221 | chr5 | q22.1 | 111247929 | 135 | CAMK4 | intron_variant | CpG: 89 | 23181 |
| Donor2_222 | chr5 | q14.1 | 81666106 | 919 | SSBP2 | intron_variant | CpG: 217 | 83873 |
| Donor2_223 | chr5 | q11.2 | 56518036 | 306 | C5orf67 | intron_variant&non_coding_transcript_variant | CpG: 59 | 36630 |
| Donor2_224 | chr5 | q35.3 | 181111555 | 963 | | upstream_gene_variant | CpG: 24 | 3097 |
| Donor2_225 | chr5 | q34 | 163912199 | 1402 | | intergenic_variant | CpG: 58 | 406259 |
| Donor2_226 | chr5 | q34 | 163846274 | 1073 | | intergenic_variant | CpG: 58 | 340334 |
| Donor2_227 | chr5 | q31.2 | 139711565 | 10 | | intron_variant&non_coding_transcript_variant | CpG: 177 | 137 |
| Donor2_228 | chr5 | q15 | 93604677 | 40 | | downstream_gene_variant | CpG: 42 | 167 |
| Donor2_229 | chr5 | p13.1 | 39187109 | 968 | FYB1 | intron_variant | CpG: 139 | 111975 |
| Donor2_230 | chr5 | q11.2 | 53842710 | 1676 | | intergenic_variant | CpG: 16 | 281670 |
| Donor2_231 | chr5 | p13.2 | 35873037 | 1084 | IL7R | intron_variant | CpG: 33 | 254800 |
| Donor2_232 | chr6 | q25.3 | 159199835 | 1511 | FNDC1 | intron_variant | CpG: 174 | 29548 |
| Donor2_233 | chr6 | q25.1 | 151408056 | 604 | RMND1 | intron_variant | CpG: 153 | 16614 |
| Donor2_234 | chr6 | q24.2 | 144531706 | 331 | UTRN | intron_variant | CpG: 187 | 244562 |
| Donor2_235 | chr6 | q24.1 | 142145819 | 1057 | NMBR | intron_variant | CpG: 28 | 1235 |
| Donor2_236 | chr6 | q22.31 | 121900863 | 564 | LOC105377979 | intron_variant&non_coding_transcript_variant | CpG: 95 | 498600 |
| Donor2_237 | chr6 | q21 | 110876642 | 2149 | AMD1 | intron_variant | CpG: 180 | 269 |
| Donor2_238 | chr6 | q16.1 | 97229464 | 539 | MMS22L | intron_variant | CpG: 53 | 53357 |
| Donor2_239 | chr6 | q15 | 90191279 | 2469 | BACH2 | intron_variant | CpG: 227 | 103798 |
| Donor2_240 | chr6 | q15 | 89681145 | 809 | MDN1 | intron_variant | CpG: 40 | 42300 |
| Donor2_241 | chr6 | p12.1 | 56751604 | 962 | DST | intron_variant | CpG: 107 | 91658 |
| Donor2_242 | chr6 | p21.1 | 45387712 | 1568 | RUNX2 | intron_variant | CpG: 91 | 9188 |
| Donor2_243 | chr6 | p21.1 | 44916023 | 1930 | SUPT3H | intron_variant | CpG: 14 | 188357 |
| Donor2_244 | chr6 | p21.2 | 37017350 | 291 | FGD2 | intron_variant | CpG: 23 | 27484 |
| Donor2_245 | chr6 | q23.3 | 137716927 | 491 | | intron_variant&non_coding_transcript_variant | CpG: 82 | 150189 |
| Donor2_246 | chr6 | q12 | 66569914 | 260 | | intergenic_variant | CpG: 21 | 474976 |
| Donor2_247 | chr6 | p21.1 | 40740261 | 37 | | intergenic_variant | CpG: 29 | 140420 |
| Donor2_248 | chr6 | p22.2 | 26253283 | 971 | H2BC9 | 3_prime_UTR_variant | CpG: 27 | 1278 |
| Donor2_249 | chr6 | p22.3 | 25078230 | 945 | CMAHP | downstream_gene_variant | CpG: 36 | 61463 |
| Donor2_250 | chr6 | p22.3 | 24888969 | 377 | RIPOR2 | intron_variant | CpG: 82 | 21430 |
| Donor2_251 | chr6 | p22.3 | 24168557 | 590 | DCDC2 | downstream_gene_variant | CpG: 23 | 42273 |
| Donor2_252 | chr6 | p22.3 | 16689083 | 1042 | ATXN1 | intron_variant | CpG: 238 | 70865 |
| Donor2_253 | chr7 | q36.1 | 152437060 | 2466 | KMT2C | upstream_gene_variant | CpG: 219 | 35 |
| Donor2_254 | chr7 | q36.1 | 151515696 | 115 | RHEB | intron_variant | CpG: 189 | 3287 |
| Donor2_255 | chr7 | q36.1 | 150823383 | 1432 | LOC105375567 | non_coding_transcript_exon_variant | CpG: 124 | 22265 |
| Donor2_256 | chr7 | q36.1 | 149625661 | 548 | ZNF767P | upstream_gene_variant | CpG: 125 | 518 |
| Donor2_257 | chr7 | q36.1 | 149144009 | 892 | ZNF398 | upstream_gene_variant | CpG: 150 | 3154 |
| Donor2_258 | chr7 | q34 | 139067601 | 18 | ZC3HAV1 | intron_variant | CpG: 88 | 31328 |
| Donor2_259 | chr7 | q32.1 | 129166397 | 274 | TSPAN33 | intron_variant | CpG: 41 | 2852 |
| Donor2_260 | chr7 | q32.1 | 128439930 | 44 | RNU7-54P | downstream_gene_variant | CpG: 60 | 15695 |
| Donor2_261 | chr7 | q31.1 | 114620591 | 351 | FOXP2 | intron_variant | CpG: 61 | 301525 |
| Donor2_262 | chr7 | q21.2 | 92606059 | 691 | CDK6 | 3_prime_UTR_variant | CpG: 77 | 15500 |
| Donor2_263 | chr7 | q11.21 | 65002873 | 471 | ERV3-1 | intron_variant | CpG: 93 | 35136 |
| Donor2_264 | chr7 | p11.2 | 55448880 | 471 | VOPP1 | intron_variant&non_coding_transcript_variant | CpG: 47 | 299 |
| Donor2_265 | chr7 | q31.2 | 115845867 | 445 | | intergenic_variant | CpG: 71 | 364450 |
| Donor2_266 | chr7 | p15.3 | 24895132 | 720 | OSBPL3 | intron_variant | CpG: 134 | 84249 |
| Donor2_267 | chr7 | p12.1 | 52817294 | 1641 | | intergenic_variant | CpG: 52 | 218289 |
| Donor2_268 | chr7 | p14.2 | 36781349 | 1785 | | intron_variant&non_coding_transcript_variant | CpG: 59 | 391152 |
| Donor2_269 | chr8 | q24.3 | 140465381 | 982 | LOC105375782 | intron_variant&non_coding_transcript_variant | CpG: 61 | 430 |
| Donor2_270 | chr8 | q21.3 | 90622276 | 533 | TMEM64 | 3_prime_UTR_variant | CpG: 159 | 22886 |
| Donor2_271 | chr8 | q21.2 | 85254556 | 1333 | CA13 | intron_variant | CpG: 39 | 8617 |
| Donor2_272 | chr8 | q13.3 | 70240644 | 468 | NCOA2 | intron_variant | CpG: 251 | 161267 |
| Donor2_273 | chr8 | q12.3 | 63026555 | 21 | GGH | intron_variant | CpG: 91 | 12009 |
| Donor2_274 | chr8 | q12.1 | 59115435 | 274 | TOX | intron_variant | CpG: 175 | 2141 |
| Donor2_275 | chr8 | q12.1 | 58554249 | 652 | SDCBP | intron_variant | CpG: 72 | 705 |
| Donor2_276 | chr8 | q11.23 | 53845726 | 826 | ATP6V1H | upstream_gene_variant | CpG: 59 | 2109 |
| Donor2_277 | chr8 | p11.22 | 38573626 | 500 | LOC105379384 | intron_variant&non_coding_transcript_variant | CpG: 52 | 77156 |
| Donor2_278 | chr8 | p21.3 | 23108787 | 738 | TNFRSF10C | intron_variant | CpG: 50 | 5373 |
| Donor2_279 | chr8 | p21.3 | 21918410 | 1060 | DOK2 | upstream_gene_variant | CpG: 114 | 922 |
| Donor2_280 | chr8 | q21.13 | 80432321 | 912 | | intron_variant&non_coding_transcript_variant | CpG: 188 | 53522 |
| Donor2_281 | chr8 | q21.13 | 75701415 | 696 | | regulatory_region_variant | CpG: 91 | 293226 |
| Donor2_282 | chr8 | p23.1 | 9240103 | 1303 | | intron_variant&non_coding_transcript_variant | CpG: 129 | 88260 |

| | | | | | | | | |
|---|---|---|---|---|---|---|---|---|
| Donor2_283 | chr9 | q34.13 | 131984687 | 1154 | MED27 | intron_variant | CpG: 33 | 94887 |
| Donor2_284 | chr9 | q31.1 | 104710186 | 1077 | LOC107987105 | intron_variant&non_coding_transcript_variant | CpG: 78 | 37440 |
| Donor2_285 | chr9 | q31.1 | 100090898 | 172 | ERP44 | intron_variant | CpG: 69 | 7596 |
| Donor2_286 | chr9 | q22.32 | 95424823 | 488 | LOC105376156 | intron_variant&non_coding_transcript_variant | CpG: 22 | 73443 |
| Donor2_287 | chr9 | q21.32 | 83982842 | 666 | RMI1 | intron_variant | CpG: 177 | 1615 |
| Donor2_288 | chr9 | q21.12 | 69739590 | 631 | PTAR1 | intron_variant | CpG: 74 | 19903 |
| Donor2_289 | chr9 | q21.33 | 85354065 | 741 | | intergenic_variant | CpG: 42 | 168630 |
| Donor2_290 | chr9 | p24.1 | 6891826 | 419 | KDM4C | intron_variant | CpG: 154 | 132954 |
| Donor2_291 | chr9 | p24.1 | 5661388 | 420 | RIC1 | intron_variant | CpG: 107 | 31645 |
| Donor2_292 | chrUn_JTFH010013 | | 1818 | 1569 | | | . | . |
| Donor2_293 | chrX | q28 | 155886763 | 1023 | VAMP7 | intron_variant | CpG: 36 | 5249 |
| Donor2_294 | chrX | q28 | 154488864 | 528 | UBL4A | upstream_gene_variant | CpG: 74 | 1452 |
| Donor2_295 | chrX | q23 | 110797793 | 586 | CHRDL1 | upstream_gene_variant | CpG: 45 | 1651 |
| Donor2_296 | chrX | q22.2 | 104187827 | 1006 | FAM199X | intron_variant | CpG: 83 | 20716 |
| Donor2_297 | chrX | q21.2 | 86824969 | 1237 | DACH2 | intron_variant | CpG: 66 | 676096 |
| Donor2_298 | chrX | p21.3 | 25017264 | 1213 | ARX | upstream_gene_variant | CpG: 28 | 442 |
| Donor2_299 | chrX | q25 | 125794935 | 387 | | intergenic_variant | CpG: 104 | 370328 |
| Donor2_300 | chrX | p22.33 | 3295534 | 440 | | intergenic_variant | CpG: 20 | 14794 |
